# Embryotoxic and teratogenic effects of polyethylene microbeads found in facial wash products in Zebrafish (*Danio rerio)* using the Fish Embryo Acute Toxicity Test

**DOI:** 10.1101/2020.09.16.299438

**Authors:** Margaret C. De Guzman, Patricia Anne P. Chua, Franceska S. Sedano

**Author notes:** Corresponding Author: (MCDG). (PAPC) (FSS). These authors contributed equally to this work.

## Abstract

Use of polyethylene beads in facial cleansers has been continuously questioned by scientific communities for they adversely affect aquatic organisms once these beads find their way into their habitats. This study specifically aims to determine *Danio rerio* mortality rate using lethal endpoints and to evaluate sublethal teratogenic effects in *Danio rerio* due to polyethylene microbead exposure. *Danio rerio,* a model organism for ecotoxicology, was subjected to the Fish Embryo Acute Toxicity Test. Embryos were exposed to polyethylene microbead suspensions (PE-MBS) of varying concentrations (i.e., 20 μg/L, 200 μg/L, 2000 μg/L). They were also exposed to 5% ethanol (positive control), reconstituted water (negative control), 0.01% Tween 80 (emulsifier control), and 1% DMSO (solvent control). Toxicological endpoints (i.e., egg coagulation, lack of somite formation, non-detachment of tail, and lack of heartbeat) were observed every 24 hours until the 96th hour exposure. Hatching was observed from 48 hpf while teratogenicity was observed at 144 hpf. Significant differences between means and variances were observed for all treatment groups in relation to the negative control. For all groups, 0.01% Tween 80, 1% DMSO and 20 μg/L PE-MBS did not significantly differ with the negative control due to negligible concentration but 5% ethanol and higher concentrations of PE-MBS did. This indicated that high concentrations of PE-MBS exposure may induce early hatching, mortality, increased malformation, and increased heart rate. Tukey Kramer *post hoc* Test substantiated that PE-MBS toxicity is dose dependent since embryotoxicity and teratogenicity increases at higher concentrations. LC_50_ obtained using probit analysis based on experimental data was 2455.096 μg/L, and was higher than the concentrations used in this study. Further studies should be conducted to know more about the adverse effects of polyethylene microbeads to the biota.

**Author Summary:** Margaret De Guzman, MSc, Patricia Chua, and Franceska Sedano have all equally contributed to this work in conceptualization, formal analysis, funding acquisition, and investigation. All authors have also equally headed project administration, procurement of resources and writing.

## Introduction

### Background of the Study

Marine pollution caused by plastic microbeads has been an emerging concern for the embryonic development and cellular health of many organisms [1]. Polyethylene microbeads (PE-MB) are polysynthetic resins found in beauty products and generally serve as abrasives or bulking agents in cleaning products and exfoliants in numerous beauty products [2]. Due to their minuscule size, most sewage treatment plants are unable to effectively filter these microbeads. As a result, these microplastics infiltrate the aquatic ecosystem and pose adverse effects to its constituents. Since microbeads are usually treated with additives and plasticizers during the production process [3], and have the ability to adsorb chemical pollutants [4], exposure to these microbeads may result in developmental toxicity in aquatic organisms [5].

Substances that may cause physical or functional defects in a developing embryo are considered to be teratogenic [6]. Polyethylene, the most common type of plastic used for microbeads [7], is a polymer of repeating CH_2_ units [8]; however, this polymer degrades for a long period of time [9], rendering them to be highly persistent and toxic to the environment. The chemical composition and ability of polyethylene to be carriers of toxins from industrial manufacturers makes it a potential teratogen to living organisms [10]. Once these microbeads come in contact with low trophic organisms such as fish larvae, exposure to toxic additives contained in polyethylene microbeads may interfere with metabolic pathways, alter gene integrity, and consequently lead to embryotoxicity and formation of teratogenic abnormalities [5,11].

Currently, there are no known studies conducted regarding the quantity of polyethylene microbeads in Philippine waters as well as the harmful effects that they pose. This study differs from others as this study is limited to polyethylene among other microplastics such as polypropylene, polyester, polyethylene terephthalate, and nylon. This study will also use polyethylene microbeads with sizes based on the ones contained in facial cleansers being commercially sold in the Philippines. This study will also be done *in vitro* instead of obtaining polyethylene bead samples from the marine or freshwater environment.

*Danio rerio* is the chosen test organism of the study. *Danio rerio* is a tropical freshwater fish and is readily available, inexpensive, exhibits high fecundity [12] and rapid development [13]. Its genes are also likened to 70% of genes in humans [14]. Furthermore, transparency of *Danio rerio* embryos allows researchers to observe teratogenesis in the embryonic development of the zebrafish [13]. In this study, *Danio rerio* were subjected to the Fish Embryo Acute Toxicity Test [15]. The Fish Embryo Acute Toxicity Test is used to evaluate the toxicity of certain chemicals on the embryonic development of vertebrates [16] and exposes fertilized eggs to varying concentrations of the toxin for 96 hours [17].

### Statement of the Problem

Do polyethylene microbeads induce embryotoxic and teratogenic effects on *Danio rerio* embryos?

### Research Objectives

The main objective of this study is to assess if polyethylene microbeads can induce teratogenic and embryotoxic effects in *Danio rerio.* The study specifically aims to:

1. to determine the mortality rate of *Danio rerio* using lethal endpoints such as lack of heartbeat, coagulation of fertilized eggs, lack of somite formation, and lack of detachment of tail-bud from yolk sac
2. to determine the lethal concentration (LC_50_) or the minimum concentration that is lethal to 50% of the exposed population, and;
3. to evaluate the sublethal teratogenic effects of polyethylene microbead exposure such as yolk sac and pericardial edema, bent tail and spine axes, and deflated swim bladder.

### Significance of the Study

This study is deemed significant as it provides information on the deleterious effects of polyethylene microbeads found in facial cleansers on the developing embryo of freshwater organisms such as *Danio rerio*. In addition, this study is timely and relevant since there has been an observed increase of microplastics in marine and freshwater environments [18], resulting in biomagnification and bioaccumulation [19]. Due to this occurrence, the Microbead-Free Waters Act has been observed in Canada, America, and the United Kingdom. According to Romero [20], this act may also be passed by Senator Loren Legarda in the Philippines soon, a country known to be third in the list of countries with the most ocean plastic pollution in a 2015 study conducted by the University of Georgia. Additionally, data gathered from this study may prompt institutions to take action in protecting bodies of water from plastic pollution and encourage local as well as international skincare manufacturers to produce a more environmentally friendly exfoliant alternative to polyethylene microbeads.

### Scope and Limitations

This study is primarily focused on the assessment of teratogenic and embryotoxic effects of virgin polyethylene microbeads in *Danio rerio*. The size of polyethylene microbeads used in the study is based on the size of commercially sold facial cleansers that contain polyethylene microbeads in the Philippines. Embryotoxic and teratogenic effects induced by polyethylene microbeads on zebrafish embryos will be assessed in accordance with the Fish Embryo Acute Toxicity Test.

### Review of Related Literature

#### Polyethylene

Polyethylene is one of the most widely manufactured polymers in the industry [21]. Its structure consists of a long chain of carbon atoms, with two hydrogen atoms attached to each carbon atom. It is a highly versatile material that can be used to make plastic bags, plastic films, bottles, and microbeads. The plastic’s melting point ranges from 110-130°C, making it highly malleable. Despite its malleability for producing a wide variety of products, it makes it a poor candidate for recycling. Despite its universal use, improper disposal of polyethylene microplastics into bodies of water makes it a vector for heavy chemical adsorption. Heavy metals such as cadmium and lead are adsorbed by these plastics and can be detrimental for both wildlife and humans [22]. It has also received criticism for containing pro-oxidants and disintegrating into smaller fragments upon exposure to light, heat, and oxygen [23].

#### Microbeads

Since its introduction to the industry in 1972, microbeads have been a popular ingredient in facial washes and facial scrubs as they serve to exfoliate and scrape away dry cells from the surface of the skin [24]. They are also incorporated in soaps and function as abrasives that remove dirt and debris found in the epidermis. The presence of plastic microbeads has been increasing in aquatic systems and yet its presence has only received attention a few years ago. Recently, microbeads have been given much attention such that countries such as Canada, New Zealand, United Kingdom, and the United States of America have banned the use of microbeads in commercial products [25]. A study conducted by Jingyi et al. [10] found that due to the continuous increase of synthetic plastic production in beauty product companies and poor management of plastic waste, water pollution by microbeads has exponentially escalated and has been a great issue of concern from public authorities. In 2018, a study found many urban areas with a maximum microplastic concentration of about 517,000 particles m**^-2^** [26]. Additionally, evidence of plastic microbeads from beauty products has been reported to bypass sewage treatments and found afloat in Hong Kong bays [27] while microplastic fragments and polyethylene microbeads mistaken for fish food were found in the gastrointestinal tract of commercial Japanese Anchovy [28]. Ingested microbeads have also affected other deep-sea organisms such as mussels and oysters. These bivalves were found to contain 0.36 to 0.47 particles of microplastic per gram [29]. Recent evidence has also shown that microplastics such as microbeads have the capacity to adsorb toxic chemicals, carry harmful bacteria and release them in digestive systems once ingested [30]. The production process of polyethylene microbeads usually include intentional treatment of additives such as flame retardants, plasticizers, pigments, and UV stabilizers as these additives prevent fire hazards and maintain product integrity [31] (Gallo et al., 2018). Polyethylene may also contain some monomers such as vinyl chloride and Bisphenol A (BPA) that contain endocrine disrupting components and induce adverse effects upon ingestion or exposure [31]. In a similar study, when mice were fed with microbeads, microplastics were seen to accumulate in the liver, kidney, and intestines. The increase of this foreign substance in bodily tissues have also heightened the levels of oxidative stress in mice [32]. In lieu of microbeads easily adsorbing pollutants, another pollutant associated with microbeads is polybrominated diphenyl ethers (PBDEs) known to aclack fontcumulate in shellfish consumed by humans. A study by Wardrop et al. [33] showed PBDE accumulation of up to 12.5% sourced from polyethylene microbeads of Nivea Exfoliating Face Scrub in the rainbow fish. PBDE pollutant is associated with neurological, fertility, and immune system problems, biomagnifying the aquatic food chain [4]. In effect, these studies have been a rising concern for humans and animals alike.

#### Facial washes with polyethylene microbeads in the Philippines

The prevalence of microplastic pollution is not uncommon in the Philippines as it ranked third in the world for the highest plastic waste inputs into the ocean [34]. Statistically, the Philippines generates about 0.28 to 0.75 million metric tons of plastic litter, yearly [34]. Studies by Kalnasa et al. [35] and Paler et al. [36] investigated the occurrences of microplastic litter in Macajalar Bay and Southwestern Luzon, respectively and revealed that a large percentage of plastic litter were brightly colored spherules. Another study by Bucol et al. [37] quantified microplastics ingested by rabbitfish (*Siganus fuscescens*) from coastal areas of Negros Oriental and found an average of 0.6 particles/fish. These microplastic spherules were speculated to have originated from facial cleansers and other cosmetic products that contain microbeads.

In the Philippines, there are a number of facial cleansers being sold in the market that contain polyethylene microbeads such as Oil-free Acne Wash Daily Scrub, Clear Pore Daily Scrub, and Deep Action Exfoliating Scrub [38]. The rise of microbead consumption and worsening of marine litter over the years have prompted government officials like Senator Loren Legarda to draft a bill that seeks to ban microbead production in the Philippines last 2018 [20] to mitigate microbead production just as New Zealand, Austria, Belgium, and the Netherlands have. In addition, Senator Loren Legarda also proposed to file a bill that will ban microplastic consumer products and single-use plastics that would otherwise bring harm to the environment [39]. In 2017, EcoWaste Coalition, along with other private groups such as Coastal Conservation, Marine Conservation Philippines, and many others endorsed a letter to the Department of Health (DOH) and Food and Drug Administration (FDA) pleading for an expedited implementation of the microbead ban. These groups stated that since plastic microbeads in drainage systems leach their way into the bodies of water, quick action must take place before they negatively affect the food chain, especially those who consume seafood [40]. Currently, DENR issued a resolution of Republic Act No. 9003 that implements the ban of single use plastics in the Philippines [41].

#### Danio rerio

In this study, *Danio rerio* was chosen as the test organism. The zebrafish is a valuable genetic model system for the study of developmental biology and disease [42]. They are prolific breeders that can lay up to hundreds of eggs per week [43], exhibiting high fecundity and rapid development. Their lifespan can reach up to 5 years and are omnivorous in nature [44]. For the past several years, the use for *Danio rerio* for scientific studies has been popular as it provides optical clarity when observing embryos with developing pathologies [45] as well as the developmental stages of a typical organism. They also have a high degree of genomic conservation and are likened to humans in terms of cellular, molecular, and physiological processes [42]. Furthermore, these genetic and physiological similarities with humans include the brain, digestive tract, musculature, vasculature, and innate immune system. 70% of human disease genes also have similar homologs found in the genes of *Danio rerio* [46]. *Danio rerio* are preferentially used for embryonic studies as they allow clear visualization of the dynamics of organogenesis using a simple stereomicroscope [42]. *Danio rerio* has been used in many toxicity studies for the reason that it is one of the best-known models of vertebrate development. The use of *Danio rerio* in studying the toxicity of microplastics is not uncommon. Despite toxicity of microplastics being common, this study differs from other studies because it focuses on polyethylene microbeads found in commercial products such as facial washes commonly used in the Philippines.

#### Embryotoxic and teratogenic effects of microplastics

Teratogens are agents or substances that induce abnormality following fetal exposure. Likewise, teratology is the study of abnormal development in embryos and the causes of congenital malformations or birth defects. These teratogens may be present on either the body surface or internal to the viscera [47]. Embryotoxicity, on the other hand, refers to injury to the embryo resulting in death or abnormal development due to exposure to toxic substances [48]. A study by Oehlmann et al. [49] conducted showed that ingestion of microplastics can affect reproduction and hormone function of marine animals like annelids, mollusks, crustaceans, insects and fish. When retained in the internal viscera for an extended amount of time, ingested microplastics may cause reproduction malfunction, increased risk of death, bioaccumulation, and even eggshell thinning [50].

However, a similar study conducted by Batel et al. [51] found that exposure to microplastics did not induce morphological effects on *Danio rerio* embryos nor did microplastics permanently accumulate in adult *Danio rerio* gills under 6 or 24 hours of incubation.

Another study stated that exposure to 1000 μg/L of microplastics significantly lessened swimming competence and speed in larval zebrafish. At gene level, this exposure resulted in upregulated expression of genes concerning “inflammation (il1b) and oxidative stress (cat)” [52]. In relation to exposure of zebrafish to microplastics, a study by Cormier et al. [53] stated that microplastics may be vectors for organic pollutants such as oxybenzone (BP3), benzo[a]pyrene (BaP), and perfluoro octane sulfonate (PFOS). This exposure effected alteration in cyp1a gene transcription, larval swimming behavior, and hatching rate at 72 hours. For other organisms such as *H. azteca,* exposure to specifically polyethylene microplastics was found to cause lessened organism growth and significant decrease of reproduction for 5000 and 10,000 microplastics/mL [54].

A study by Gallo et al. [31] stated that exposure of marine organisms to micro and nano plastics results in bioaccumulation and adverse toxic endpoints as these microplastics contain endocrine disrupting properties such as alkylphenols, bisphenol A (BPA), and phthalate esters (DEHP) in concentrations as high as 500,000 mg/kg (ppm). The presence of microplastics increases BPA uptake in *Danio rerio* and in turn causes gene-upregulation in the central nervous system and inhibition of acetylcholinesterase (AChE), which entail that microplastics are neurotoxic [55]. A similar study by Nobre et al. [56] studied the effects of polyethylene pellets on the embryonic development of *Lytechinus variegatus* (sea urchin) and found that exposure to these microplastics induced toxic effects and increased anomalous embryonic development by 58.1% and 66.5% respectively. These findings substantiate that plastic pellets have the ability to act as vectors of pollutants that include additives contained in the surface of virgin pre-processed particles.

Other studies show the detrimental effects of microbeads to aquatic organisms, particularly in *Danio rerio*. A study by Träber et al. [57] implanted polyacrylamide beads into developing *Danio rerio* embryos to quantify cell-scale stress in its morphogenesis and organ formation. Stresses induced by microbead implantation had a detrimental effect on neural rod formation.

Lei et al. [58] observed that microplastics such as polyamides, polyethylene, polypropylene, polyvinyl chloride, and polystyrene cause intestinal damage and other adverse effects in zebrafish and in the nematode, *Caenorhabditis elegans* in freshwater pelagic and benthic environments. Absence or insignificant levels of lethality were observed in the zebrafish upon exposure at 0.001-10 mg L^-1^ microplastics for 10 days. Meanwhile, concentrations of microplastics at approximately 70 μm resulted in intestinal damage, cracking of villi, and splitting of enterocytes in zebrafish. On the other hand, exposure of 5.0 mg m^-2^ microplastics for 2 days notably impeded survival rates, reproduction and body length of the nematodes. For both organisms, exposure to microplastics at specific sizes contributed to decrease in calcium levels and increased expression of the glutathione S-transferase 4 enzyme in the intestine. This increase confirmed intestinal damage and increase of oxidative stress as effects of exposure to specific concentrations of microplastics. From these results, researchers suggested that toxicity of microplastics were based on size instead of their composition [58]. Although past toxicity studies related to polyethylene microplastics already exist in literature, this study differs since it investigated embryotoxicity and teratogenicity of polyethylene microbeads based on sizes found in commercially sold facial cleaners in the Philippines.

#### Fish Embryo Acute Toxicity Test

The Fish Embryo Acute Toxicity Test (FET) is a method used to study chemical toxicity in aquatic ecosystems in vivo [59]. This test (FET) is deemed advantageous for studies that need to observe the fish under varying concentrations of the test solution. Fish is primarily used in toxicity testing because of their metabolic capacities and they are, more often than not, the primary targets of water pollution and heavy metal effluents [12]. A study by Gülden et al. [60] compared cytotoxicity data from fish and mammalian cell lines and found that both are equally sensitive. This study produced evidence that the Fish Embryo Acute Toxicity Test, while it is performed on fish, is extremely relevant in humans as they both show high sensitivity to FET tests. It uses 4 toxicological endpoints for the determination of toxicity in zebrafish eggs: coagulation of fertilized egg, lack of somite formation, detachment of the tail-bud from yolk sac, and lack of heartbeat [16].

### Methodology

#### Danio rerio maintenance

Thirty (30) female and twenty-five (25) male *Danio rerio* approximately 5-months-old and void of any pharmaceutical treatment were purchased from Freshwater Aquaculture Center, College of Fisheries in Central Luzon State University Science City of Munoz, Nueva Ecija. Female and male *Danio rerio* were separated and placed in two 15-gallon glass tanks three fourths (¾) filled with dechlorinated water that was maintained at 26 ± 1 °C, well-aerated with dissolved oxygen at a concentration of 6.6 mg/L, electrical conductivity of 0.256 mS/cm, water hardness of 185 mg/L CaCO_3_ and at a constant pH of 7.2± 1 [17]. These conditions were maintained with the use of API Freshwater Master Test kit. The feeding regime consisted of *Danio rerio* being fed with 300 µm of Tetra®Min Tropical Flakes twice a day at 8:00 am and 4:00 pm daily. This slightly deviated from the original OECD protocol of requiring to feed *Danio rerio* with dry flake food and brine shrimp 3-5 times daily. *Danio rerio* were subjected to a 12-hour-light cycle and were acclimated for two weeks prior to the experiment. The fish were fed with egg yolk the night before mating to increase the likelihood of breeding [61].

#### Preparation of polyethylene microbead suspensions

Clear polyethylene microbeads (PE-MB) 300-355 μm in diameter and 1.10 g/cc in density were purchased from Copsheric LLC (Santa Barbara, CA). These measurements were chosen based on a study of Chang [38] who characterized microbeads from various commercial facial exfoliating cleansers. Since PE-MB are hydrophobic in nature, they were treated with 0.01% Tween 80, a surfactant used to disperse hydrophobic particles in aqueous solutions. To prepare 0.01% Tween 80 solution, a beaker was filled with distilled water and was brought to a boil for 5 minutes. 0.1 g of Tween 80 per 100 ml was slowly dispensed in boiled water. After cooling, the desired amount of PE-MB was added to a test tube and was placed in a vortex mixer for at least five minutes. PE-MB was left to soak in 0.01% Tween 80 for 24 hours or until equal dispersion was achieved [62]. Polyethylene microbeads were then filtered from 0.01% Tween 80 using Whatman® Grade 1 filter paper with a pore size of 11 μm. Afterwards, polyethylene microbead suspensions (PE-MBS) were prepared by adding polyethylene microbeads to a solution consisting of reconstituted water and 1% DMSO, an organic solvent capable of softly dissolving PE-MB and producing a suitably concentrated stock solution [17]. Sterile and aerated reconstituted water was used in the preparation of PE-MBS [63]. Three concentrations of PE-MBS were prepared based on previous studies on microplastic toxicity by [63] that used the same concentrations [1]. The three concentrations of microbead test suspensions used in this study are 20 μg/L, 200 μg/L and 2000 μg/L.

#### Egg production and collection of fertilized eggs

*Danio rerio* eggs were collected using mass spawning. The number of *Danio rerio* used for mass spawning from the original OECD protocol was modified to acquire the desired number of *Danio rerio* eggs for the study. Groups of *Danio rerio* with a sex ratio of 1 female: 3 males were placed in spawning tanks [64] and were exposed to a 14-hour-light cycle the day before the eggs were collected [17].

A spawn trap was placed inside the spawning tank as a means of collecting *Danio rerio* eggs. Spawn traps were covered with an inert wire mesh with a size approximately 2±0.5 mm to prevent predation by adult *Danio rerio*. Mating, spawning and fertilization took 30 minutes after the onset of light on the day of testing and eggs collected through spawn traps [17]. After collecting the eggs from the spawning tank, they were rinsed with reconstituted water. Reconstituted water consisted of 294.0 mg/L CaCl_2_·2H_2_O, 123.3 mg/L MgSO_4_·7H_2_O, 63.0 mg/L NaHCO_3_, 5.5 mg/L KCl [65]. A volume of 0.05 ml of Methylene blue was also added to reconstituted water to prevent fungal and parasitic infection that may occur in *Danio rerio* eggs [66]. The reconstituted water solution was aerated for a minimum of 24 hours before being used in the experiment. Fertilized eggs were sorted from unfertilized eggs. The fertilized eggs were transferred to multi-well plates with the reconstituted water. The number of unfertilized and fertilized eggs were counted to check the validity of the results obtained from the Fish Embryo Acute Toxicity test as overall fertilization rate of all eggs collected must be ≥ 70% in the batch tested [17].

### Fish Embryo Acute Toxicity Test

The fertilized eggs were immersed in the test solution immediately after egg collection. The viable fertilized eggs and the unfertilized eggs were separated and counted for raw data. After separation, 60 viable fertilized eggs per treatment group were placed in a chamber containing their respective test concentrations (Table 1) for initial exposure [17]. A dropper was used to transfer viable fertilized eggs from their respective chambers to 96–well plates containing microbead test suspensions. For the experimental set-up of the Fish Embryo Acute Toxicity Test, 20 *Danio rerio* embryos per test concentration with 3 replicates each will be placed in 96-well plates containing the test concentrations.

**Table 1.** Experimental setup showing the composition and volume of each treatment used in the Fish Embryo Acute Toxicity test.

| Treatment | Volume in mL | Composition |
| --- | --- | --- |
| Negative control | 0.5 | Reconstituted Water |
| Internal Plate Control | 0.5 | Reconstituted Water |
| Positive Control | 0.5 | 5% ethanol |
| Solvent Control | 0.5 | 1% DMSO |
| Emulsifier Control | 0.5 | 0.01% Tween 80 |
| Treatment 1 | 0.5 | 20 µg/L PE-MBS |
| Treatment 2 | 0.5 | 200 µg/L PE-MBS |
| Treatment 3 | 0.5 | 2000 µg/L PE-MBS |

Shown in Table 1 is the experimental setup done for the Fish Embryo Acute Toxicity Test. Five internal plate controls containing sterile reconstituted water will be added to each 96-well plate to identify any potential contamination of the plates by the manufacturer that may be suspected to affect the outcome of the results [17]. If more than one embryo dies per plate in the internal plate control, the test is considered invalid and must be performed again. Reconstituted water was specifically used as a reference solution for negative and internal plate control as *Danio rerio* embryos have stricter requirements than adult fish and may be more susceptible to disease if incubated in regular distilled water [67]. Five percent (5%) ethanol served as the reference substance for positive control as it is known to be a neurotoxicant that induces deformations and mortality in *Danio rerio* [68]. 1% DMSO and 0.01% Tween 80 served as solvent and emulsifier controls, respectively, as they were used in the preparation of PE-MBS and to ensure that these substances do not cause embryotoxicity and teratogenic abnormalities to the organisms under investigation.

Water temperature was made sure to be maintained at 26 ± 1 °C in the test chambers at any time during the test. Certain parameters of newly fertilized *Danio rerio* eggs were checked to ensure that it is valid for the Fish Embryo Acute Toxicity Test.

The following factors were observed in the collected eggs for the test results to be valid:

1. Overall fertilization rate of all eggs collected must be ≥ 70% in the batch tested.
2. Overall survival of embryos in the negative and solvent control must be ≥ 90% at the end of the 96 hour exposure.
3. Exposure to the positive control must result in mortality not less than 30% at the end of the 96 hour exposure.
4. Hatching rate in the negative and solvent control must be ≥ 80% at the end of the 96 hour exposure.

### Observations for the Fish Embryo Acute Toxicity Test

The following toxicological endpoints were observed using a Leica ES2 Stereoscope with a magnification of 100x: (1) coagulation of embryos, (2) lack of somite formation, (3) non-detachment of the tail, (4) lack of heartbeat, and (5) hatching rate. For embryo coagulation, it was observed as milky white, yet it appeared dark under the microscope. For lack of somite formation, it should be noted that a zebrafish embryo undergoing normal development at 26 ± 1 °C, will form approximately 20 somites after a day. In addition, side-to-side contractions of the embryo signifying somite formation were observed. Lack of somite formation was also recorded after 24, 48, 72, and 96 hours. Non-detachment of the tail means absence of a posterior extension of the body of the embryo. Absence of this was recorded after 24, 48, 72, and 96 hours. Lack of heartbeat was recorded after 48, 72, and 96 hours since visibility of heartbeat occurs after 48 hours of a normally developing zebrafish embryo at 26±1°C. It should be noted that erratic heartbeat and visible heartbeat in the absence of circulation in aorta abdominalis are non-lethal. Hatching, despite not being a teratogenic endpoint involved in the calculation for LC_50_, was observed and recorded after 47, 72 and 96 hours for it ensures exposure of the embryo in the absence of a potential barrier function of the chorion.

Any positive results observed for any of the toxicological endpoints rendered the *Danio rerio* embryo dead. Moreover, hatching and heartbeat were observed in control and treatment groups from 48 up to 96 hpf were recorded as well. The remaining toxicological endpoints were recorded every 24 hours until the end of the 96 hour exposure.

At 144 hpf, *Danio rerio* larvae were euthanized using hypothermic shock. The fish were quickly immersed in an ice bath consisting of 5 parts ice and 1 part distilled water for 40 minutes or until cessation of gill and heart movement was observed [69]. Once movement was no longer visible, *Danio rerio* were mounted in glass slides with 10% glycerol. Prepared microscope slides were then observed under Leica ES2 Stereoscope with a magnification of 100x to assess the different teratogenic effects induced by PE-MBS such as yolk sac edema, pericardial edema, bent body axes, tail curvature and collapsed swim bladder.

### Statistical Analysis for Fish Embryo Acute Toxicity Test

Cumulative mortality, cumulative hatching, number of malformations, and the number of embryos that represent coagulation, lack of somite formation, non-detachment of tail, lack of heartbeat, and hatching, respectively for all treatments after the 24, 48, 72, 96 hour exposure were recorded. Probit analysis for the estimation of LC**_50_** values at 96 hour exposure for mortality with a 95% confidence limit was recorded for graphing and interpretation as well [17]. It should be noted that LC**_50_**

Treatment effects of the different concentrations of microbead suspensions on the developmental parameters and mortality of *Danio rerio* embryos were determined using one-way analysis of variance (ANOVA). Kruskal-Wallis test was performed if data did not pass Shapiro-Wilk’s test of normality. Dunnett’s test was used to compare the treatment means with their corresponding controls if parameter assumptions of normality and homogeneity of variances were met whereas Dunn’s test was used to analyze obtained data if assumptions were not met. Multiple comparisons among the three treatments were performed through Tukey-Kramer *post hoc* test. Statistical analyses were executed using Microsoft Excel Real Statistics Software. Data is significant for p ≤0.05.

### Institutional Animal Care and Use Committee (IACUC)

Under guidelines of the Institutional Animal Care and Use Committee, *Danio rerio* was used in this study for the interest of relevance to human and animal health, to improvement of knowledge, and to the good of society [70]. Researchers involved in the experiment ensured proper handling of the specimens.

Factors affecting the housing and feeding of the *Danio rerio* such as UV-sterilization, ventilation, aeration (6.6 mg/L O_2_) temperature (26 ± 1 °C), water cleaning, water salinity (185 mg/L CaCO_3_), electrical conductivity (0.256 mS/cm), and pH (7.2 ± 1) were adjusted in accordance to the proper care and breeding of *Danio rerio* as earlier mentioned in the methodology.

*Danio rerio* were placed and maintained in 2 15-gallon glass tanks. Dimensions of each glass tank were 20″ x 10″ x 12″. 15% of the water in glass tanks was replaced every week. Before replacement, the tap water acquired in the sanitized gallon-sized bucket was pre-treated first with a water conditioner to adjust the pH level and to remove toxins and metal residue in the tap water. Water was UV-sterilized with a portable UV water sterilizer submerged and stirred in the bucket until light of the sterilizer turned off.

After UV sterilization, the water in the tank was removed with a siphon tip placed into the tank’s substrate at the bottom. The siphon removed debris and the tank water. Temperature of the remaining water in the tank and that of the new water in the buckets were measured with a thermometer to know if temperatures were near to one another. Pretreated water was poured slowly into the tanks [71].

They were fed properly and regularly. The fish were fed twice everyday as earlier stated in the methodology.

After conducting the study, adult *Danio rerio* used for breeding were returned to their original tanks. *Danio rerio* embryos and larvae used in the Fish Embryo Acute Toxicity Test were placed in sealed plastic bags for garbage collection.

## Results and Discussion

The use of the Fish Embryo Acute Toxicity Test in this study has shown that polyethylene microbeads found in facial wash products are embryotoxic and teratogenic to *Danio rerio* embryos. Three concentrations of white PE-MBS 300-355 μm in diameter and 1.10 g/cc density were used in this study (i.e., 20, 200 & 2000 μg/L) as toxicants.

### Embryotoxicity

Polyethylene microbead embryotoxicity was evaluated using the four toxicological endpoints namely coagulation of eggs, lack of somite formation, non-detachment of tail, and lack of heartbeat [17]. Coagulated embryos are described as milky white eggs void of any structure. Lack of somite formation is characterized by absence of somites and side to side contractions. Non-detachment of tail is the inability of the embryo to extend its posterior extension while lack of heartbeat is absence of a visible heartbeat in a normally developing embryo starting at 48 hpf. Once a single toxicological endpoint is observed within the 96 hour exposure, the embryo was considered dead [17].

Cumulative mortality was observed until the 96th hour of the final static exposure. Ninety-six (96) hours were allotted for observing cumulative mortality since there are some chemicals (i.e. cationic polymers) that may not manifest their toxic potential until the embryo has been completely liberated from the protective outer shell, the chorion. In extending the static exposure to 96 hours, zebrafish development may encompass hatching [72] and cumulative mortality may be recorded as well.

Based on the results shown in Fig 1, the number of observed deceased *Danio rerio* for all control treatments and PE-MBS concentrations were in the following decreasing order: 2000 μg/L, 5% ethanol, 200 μg/L, 20 μg/L, 1% DMSO, 0.01% Tween 80, and reconstituted water for the negative control and internal plate control (S4 Appendix). The trend in Fig 1 shows that exposure to increasing concentrations of PE-MBS increased incidences of mortality as well. Upon statistical analysis, ANOVA (S9 Appendix) indicated that there is a significant difference between the means and variances of cumulative mortality of *Danio rerio* within the 96 hour exposure for all treatments.

**Fig 1.**
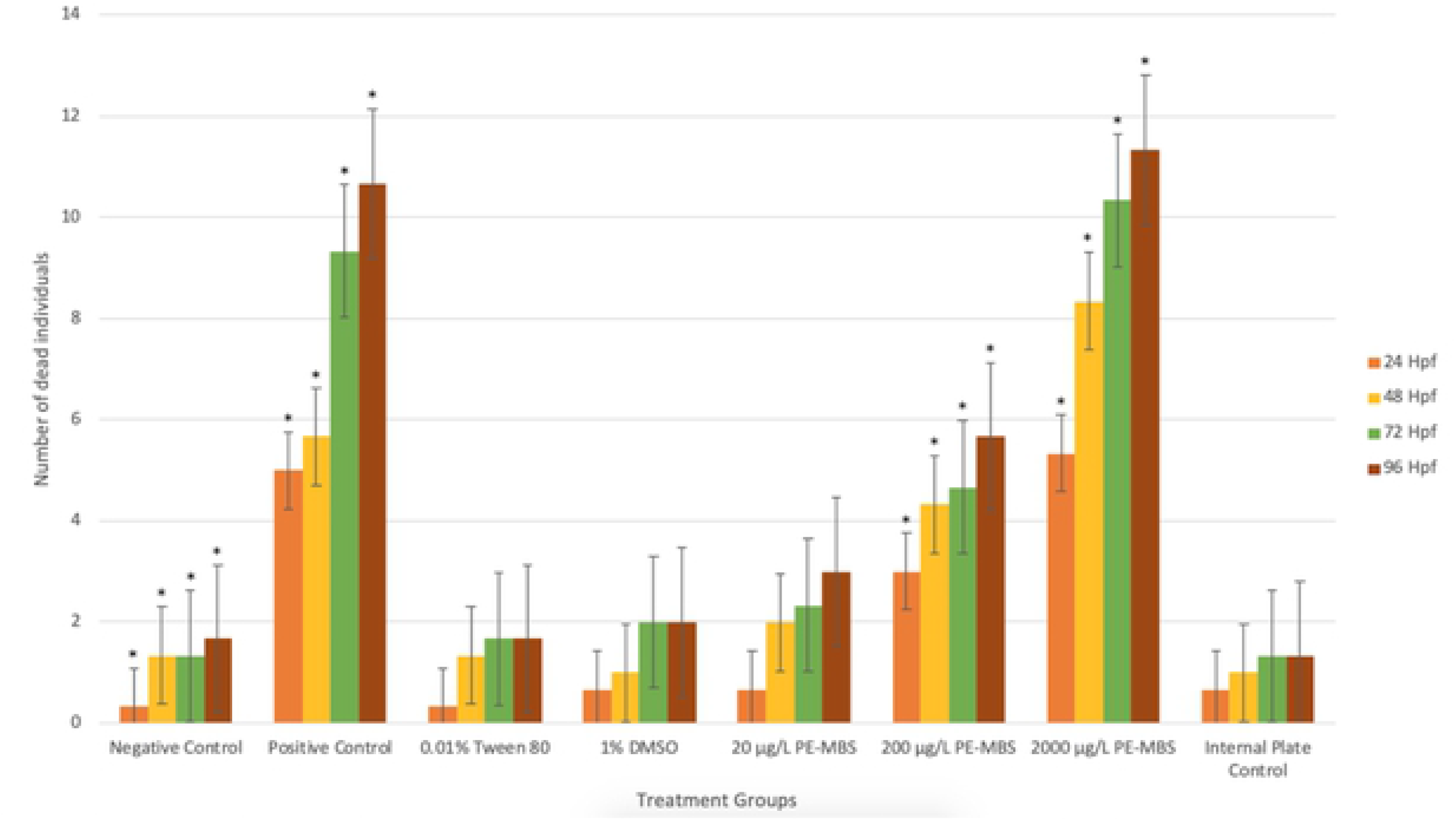
Lethal effects of PE-MBS on *Danio rerio* embryos within 96 hour exposure to different concentrations. Data shown is based on the average of three replicates performed in the study. Error bars indicate standard error. Single-asterisk indicates a statistically significant difference of cumulative mortality between *Danio rerio* (p < 0.05). (*:p < 0.05).

Based on the results found in Fig 2, coagulation accounted for the most frequently occurring lethal endpoint in all control solutions and PE-MBS concentrations, with mean percentages ranging from 38% to 80%. Lack of observable heartbeat was the second most recorded toxicological, garnering values from 0% to 33% followed by non-detachment of tail with results ranging from 0% to 29%. Lack of somite formation accounted for the least occurring endpoint in all control solutions and PE-MBS concentrations, with percent values from 0% to 12%.

**Fig 2.**
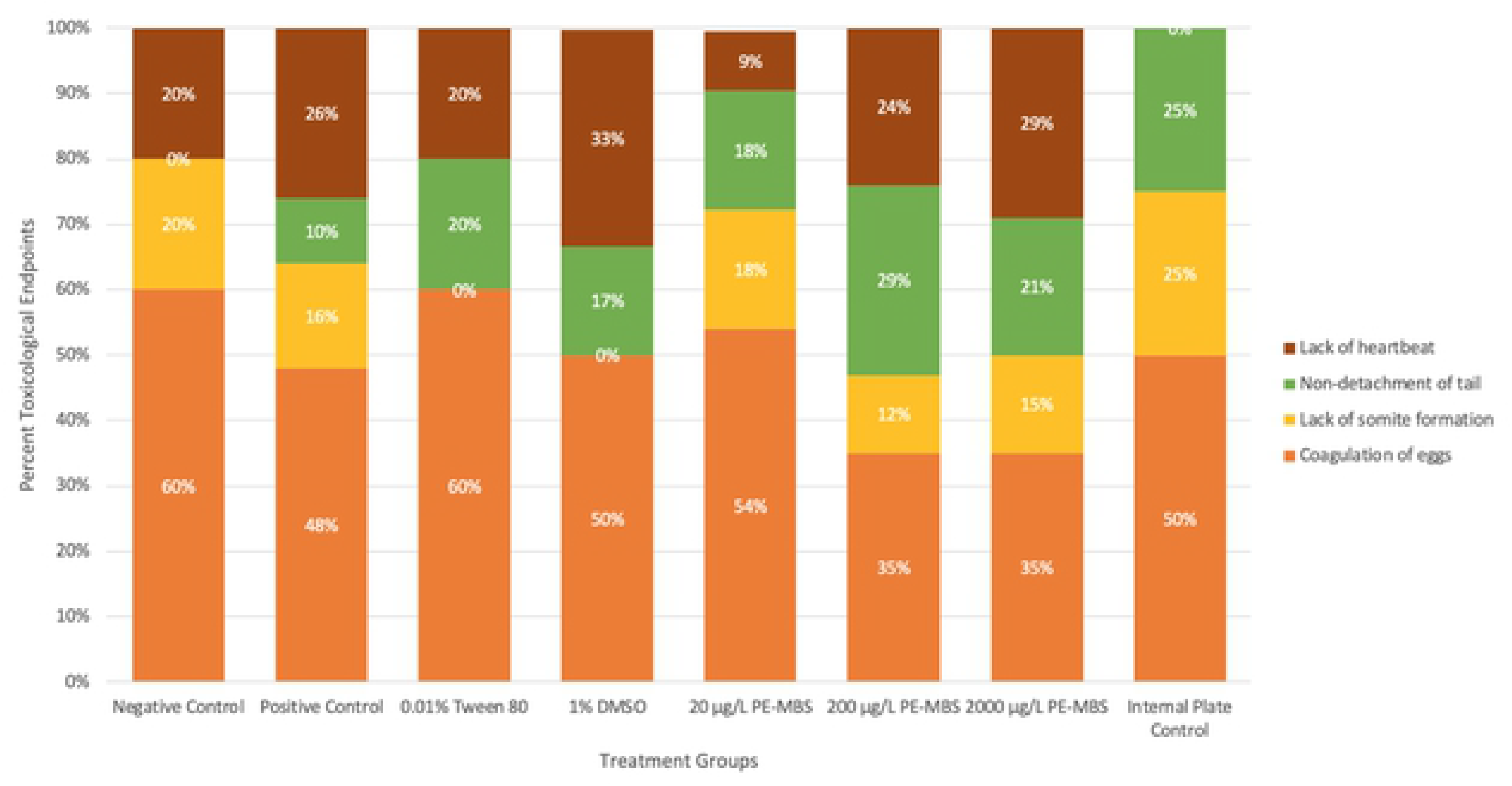
Relative percentages of toxicological endpoints observed in deceased *Danio rerio* at 96 hpf. Percentage shown is based on the average of three replicates performed in the study.

Negative controls are important since they are used to detect confounding variables [73] and serve as a basis of comparison for different test groups. Internal plate controls, on the other hand, are used to identify any potential contamination found in 96-well plates that may affect the outcome of the results [17]. Small discrepancies such as toxic endpoints observed in the negative and internal plate control may be due to extraneous variables (i.e. varying oxygen levels in well-plate, change of pressure of pipette tip into well, position of embryo in well). While extraneous variables have a wide scope that include situational variables, participant variables, investigator effects and demand characteristics; environmental factors, on the other hand, are more specific but may still fall under extraneous variables [74]. Examples of environmental variables are noise, temperature and lighting conditions of the experimental set-up.

In lieu of these toxic endpoints seen in the negative and internal plate controls, these results also coincided with toxicological studies of the zebrafish [75,76]. These studies observed low zebrafish embryo mortality in their negative controls such as dilution water [75] and buffer and egg water [76]. The latter stated that “spontaneous mortality” in the first 24 hpf may have been the reason for the mortality observed in their controls, coinciding with other literature. This may also explain the mortality of the control group in this study. Despite the mortality observed in these controls, these results had no significant difference. Additionally, requirements by OECD standards [17] stating that these controls should observe survival of at least 90% until the 96th hour were still met in this study.

Upon analysis of data gathered in the study, mortality of *Danio rerio* treated with 5% ethanol garnered results significantly different from the negative control all throughout the 96 hour exposure (S10 Appendix). Coagulation, lack of somite formation, non-detachment of tail, and lack of heartbeat were observed in deceased *Danio rerio,* with coagulation accounting for the most frequently occurring toxic endpoint. Although the mechanism behind egg coagulation remains unclear due to lack of related literature, coagulation induced by toxicant exposure is suspected to be a result of *Danio rerio* having temporal expression or lack of specific metabolic enzymes that may not allow it to metabolize harmful products during the entirety of the first 48 hours of development [77]. Exposure to toxic products such as ethanol may lead to complete cell and biomolecule disintegration as well as disruption to cell fate determination during organogenesis [78], which is manifested by milky white egg coagulation in *Danio rerio* embryos (Fig. 3C). Coagulation induced by 5% ethanol may be due to its toxic properties and ability to act as a desiccant and protein denaturant at high concentrations [79]. Lack of visible heartbeat also occurred and may possibly have been a result of its disruption of the central nervous system and inhibition of acetylcholinesterase [80] that may have caused complications related to heart failure. Data obtained from this study coincide with the study conducted by Hallare et al. [81].

**Fig 3.**
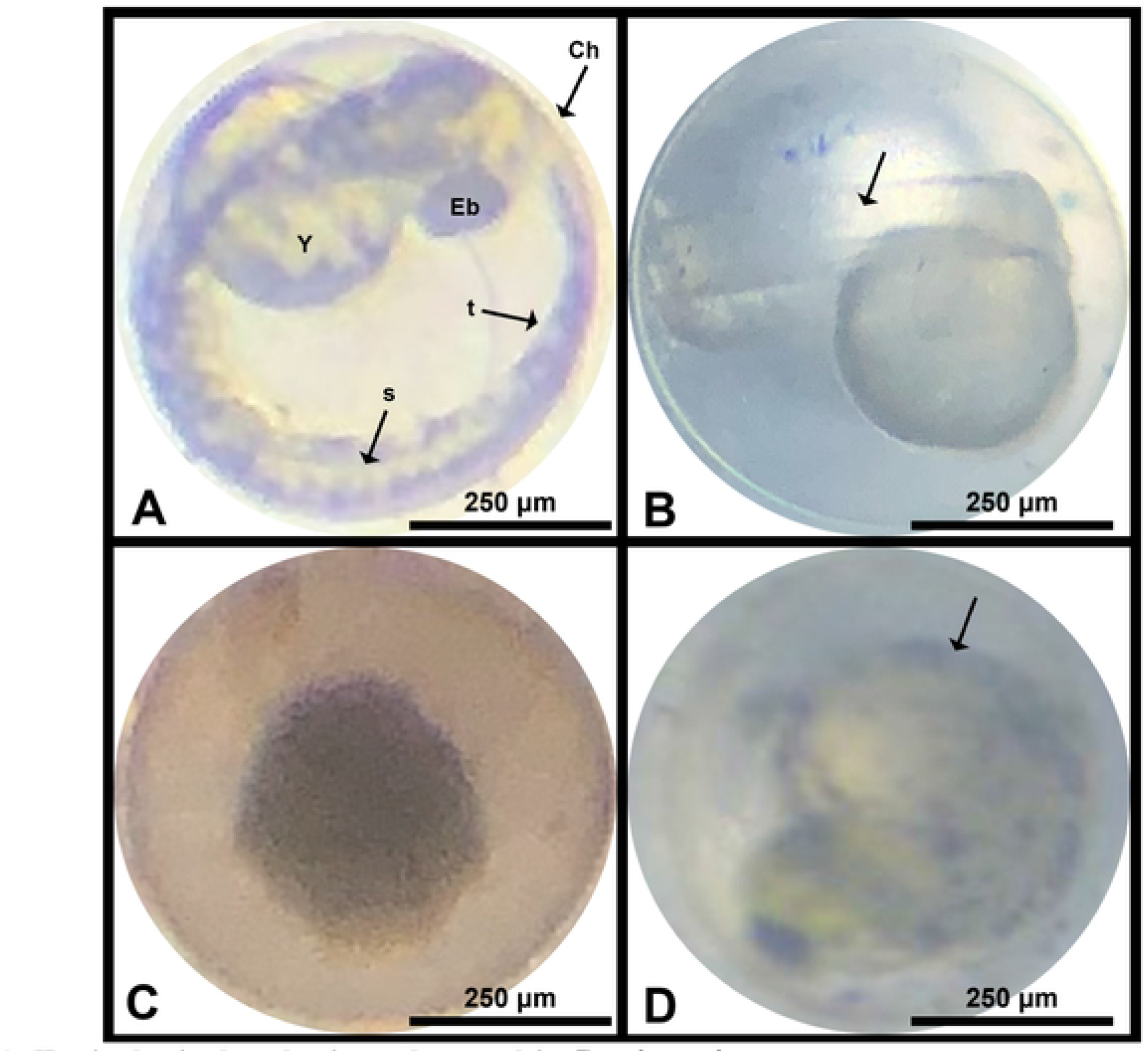
Toxicological endpoints observed in *Danio rerio.* (A) normal development of *Danio rerio* at 48 hpf observed in the negative control (RW), 0.01% Tween 80, 80% DMSO, and 20 μg/L PE-MBS. A. Embryo demonstrates eye bud (Eb), chorion (Ch), yolk (y), somites (s), and tail (t). 3 of the 4 toxicological endpoints denoting mortality: (B) lack of somite formation (*arrow*), (C) coagulation of eggs, and (D) non-detachment of tail (*arrow*) observed primarily in the positive control (5% ethanol), 200 μg/L PE-MBS, and 2000 μg/L PE-MBS.

Dunnet’s test results revealed that there is no significant difference between the means and variances of mortality obtained for 0.01% Tween 80 and 1% DMSO with the negative control all throughout the 96 hour exposure (S10 Appendix). Manifestations of cardiac failure in embryos treated with 0.01% Tween 80 may be due to its low order toxicity [82] and capability to cause electrophysiologic changes to the cardiac conduction system [83] whereas visible heartbeat observed in *Danio rerio* treated with 1% DMSO may be due to its disruption of the central nervous system and inhibition of acetylcholinesterase [80]. Although both substances are embryotoxic and inducers of various developmental effects at high concentrations as evidenced in previous studies [82,81], Tween 80 and DMSO were diluted to concentrations 0.01% and 1% respectively. Dilution of these substances were effective in making them appropriate surfactants and solvents for PE-MB [17] without causing embryotoxicity of remarkable difference with the negative control.

The results obtained for *Danio rerio* embryos treated with 20 μg/L PE-MBS did not show a significant difference with the negative control all throughout the 96 hour exposure (S10 Appendix) which suggests that 20 μg/L PE-MBS is a concentration not sufficient enough to induce embryotoxic effects to *Danio rerio* embryos. However, Dunnet’s test revealed significant differences between the means of the cumulative number of deceased *Danio rerio* treated with 5% ethanol, 200 μg/L PE-MBS, and 2000 μg/L PE-MBS with the negative control at all exposure times within the 96 hour period (S10 Appendix).

Occurrences of egg coagulation (Fig. 3C) were most frequently observed during the first 48 hours of exposure for embryos treated with PE-MBS; thus, it is speculated that coagulation is associated with a defect in the early embryonic stages of development (e.g., blastulation and gastrulation). These developmental processes are highly conserved as few alterations may cause lethal effects to the embryo [84]. Coagulation induced by PE-MBS is due to the toxic chemical components of polyethylene. According to a study by Gallo et al. [31], polymers of microplastics, even in extremely low concentrations, contain toxic chemical additives such as flame retardants, plasticizers, UV stabilizers and pigments that are intentionally treated to the surfaces of virgin polyethylene microplastics during the production process to reduce fire hazards and maintain product integrity. Another study by Rochman et al. [3] stated that virgin pre-production polyethene microplastics contain Endocrine-Disrupting Chemicals (EDCs) such as bisphenol A (BPA). Aside from BPA being an exogenous compound that interferes with metabolic pathways and proper functioning of the endocrine system [5,85], accumulated evidence from past studies have ascertained that BPA is cytotoxic, have the ability to alter gene integrity [85,86] and induce cell apoptosis and organ necrosis to developing vertebrates [87]. In this study, 1% DMSO was used as a solubilizing agent to produce a suitable suspension for polyethylene microbeads [17]. Soft extraction of polyethylene microbeads by DMSO may have caused leaching of toxic additives and other EDCs that permeated through the chorion pores, caused cell disintegration and ultimately led to incidences of milky white embryo coagulation (S2 Appendix).

Development of the heart begins at 16 hpf in which cardiac precursor cells start to differentiate and travel towards the central midline of *Danio rerio* embryos [83]. Occurrences of lack of observable heartbeat were observed in *Danio rerio* treated with PE-MBS during the embryonic and larval stage (S4 Appendix). The number of deceased *Danio rerio* due to lack of heartbeat increased upon increase in PE-MBS concentration (S2 Appendix). This may be attributed to hypoxia caused by PE-MBS exposure during the earlier developmental stages of *Danio rerio*. Accumulated evidence supports a study by Malafaia et al. [5] that polyethylene microbeads cause hypoxia in *Danio rerio* embryos as these microplastics may adhere to the chorionic membrane [88]. Since chorionic pores measure approximately less than 1 μm in diameter, polyethylene microbeads that measure 300-355 μm most likely became a barrier that hindered the passage of diffusing oxygen, and consequently interfered with gas exchange. Disruption in gas exchange results in critically low oxygen availability that induces a reactive response in which certain respiratory processes are accelerated [89] and cases of premature hatching may occur [5]. A significant number of early hatching (i.e., hatching at the 48 hpf mark) in *Danio rerio* treated with 200 and 2000 μg/L PE-MBS were observed (S5 Appendix) and may be suspected to be due to the breakdown of the chorion as a means to increase oxygen uptake in *Danio rerio* [5]. Despite greater oxygen uptake, premature hatching produces underdeveloped larvae with teratogenic abnormalities that synergistically contribute to post-hatching mortality [5]. In a study by Kuiper et al. [90], they also found that exposure of *Danio rerio* to plastic additives such as flame retardants found in microplastics contain toxic chemicals that cause high post-hatching mortality and pericardial fluid accumulation in juvenile larvae evidenced by manifestations of pericardial edema (S7 Appendix), both in which coincide with the results obtained in the conducted study.

The first sign of somite differentiation occurs after gastrulation [84]. It is from these somites that muscle cells are derived from. It is also during somitogenesis when the tail begins to extend and separate itself from the yolk. Alterations in somitogenesis due to substance toxicity affect normal development and may cause a defect in somite formation and tail detachment [84]. Incidences of embryos exhibiting lack of somite formation and non-detachment of tail were observed in *Danio rerio* treated with PE-MBS (S2 Appendix) and may be attributed to chemical additives and EDCs added to pre-production polyethene microplastics [85]. Related literature outside of this study suggests that observed toxic endpoints related to somite defect may be associated with ectodermal implications during somitogenesis [91]. Somite formation is initiated by the motion waves of gene expression that originate from the head [92], and since EDCs alter gene integrity, they may have affected normal development as well. However, further studies must be conducted to investigate to know the specific genes and the level of gene expression EDCs pose an effect on.

The concentration-mortality curve of *Danio rerio* at 96 hpf as shown in Fig 4 indicates that there is an increasing trend in mortality rate as the concentration of PE-MBS increases. Results from Tukey Kramer’s *post hoc* test (S11 Appendix) revealed that there is a significant difference between the means of the cumulative number of deceased *Danio rerio* treated with different concentrations of PE-MBS at 96 hpf which substantiates early speculations that PE-MB toxicity is dose dependent and causes concentration-dependent reduction in *Danio rerio* survival.

**Fig 4.**
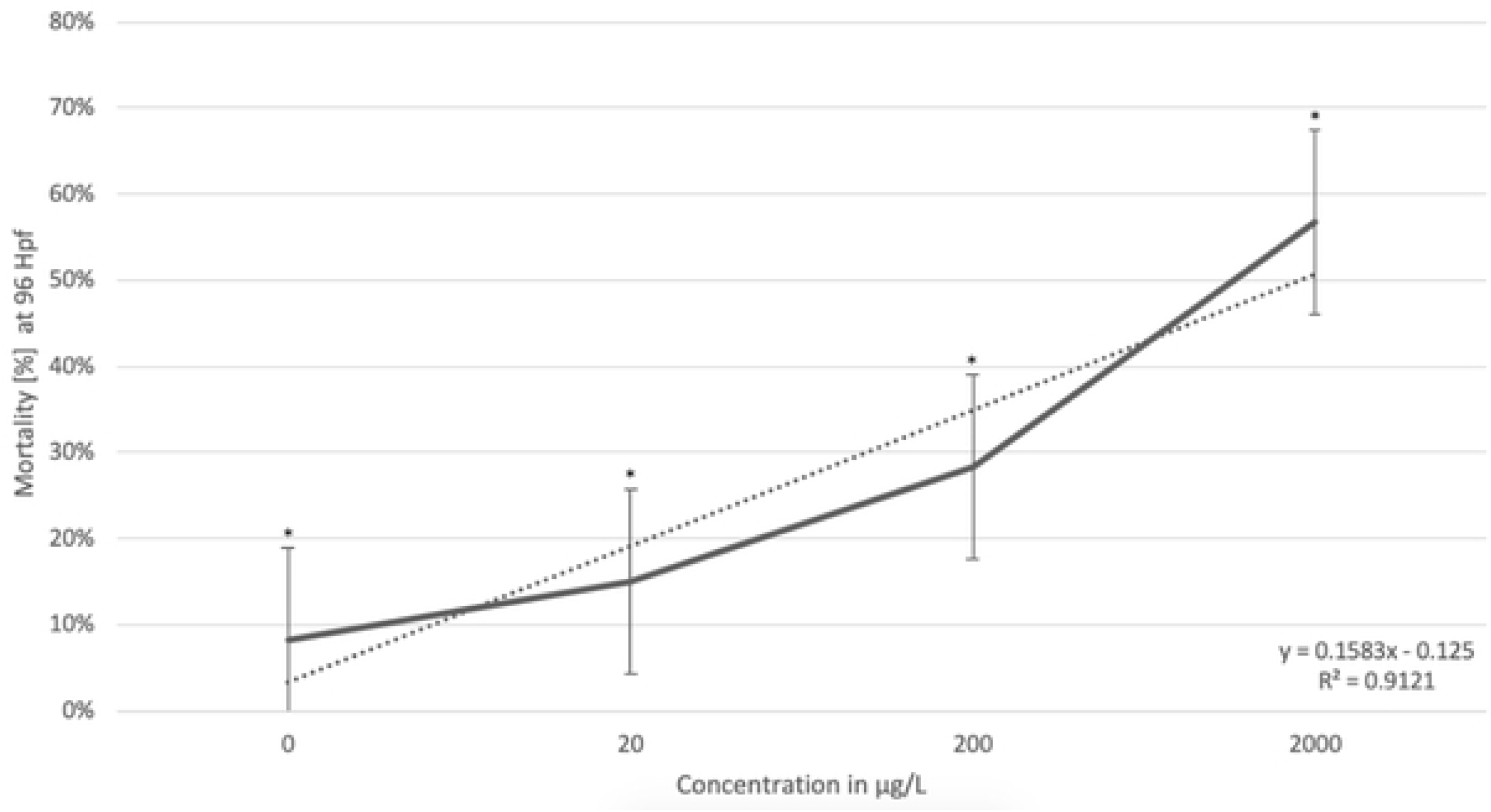
Concentration-Mortality curve in FET of *Danio rerio* treated with PE-MBS at 96 hpf. Error bars indicate standard error. Single-asterisk indicates a statistically significant difference of cumulative mortality between *Danio rerio* at 96 hpf (p < 0.05). (*:p< 0.05).

In Fig 5, the computation of LC_50_ using probit analysis, while taking into consideration results obtained from the negative control, revealed that the lethal concentration of polyethylene microbeads causing mortality to 50% of the population under study is 2455.096 μg/L (S12 Appendix). The value garnered for the LC_50_ of PE-MB is higher than treatment concentrations used in the study (i.e., 20, 200, and 2000 μg/L PE-MBS).

**Fig 5.**
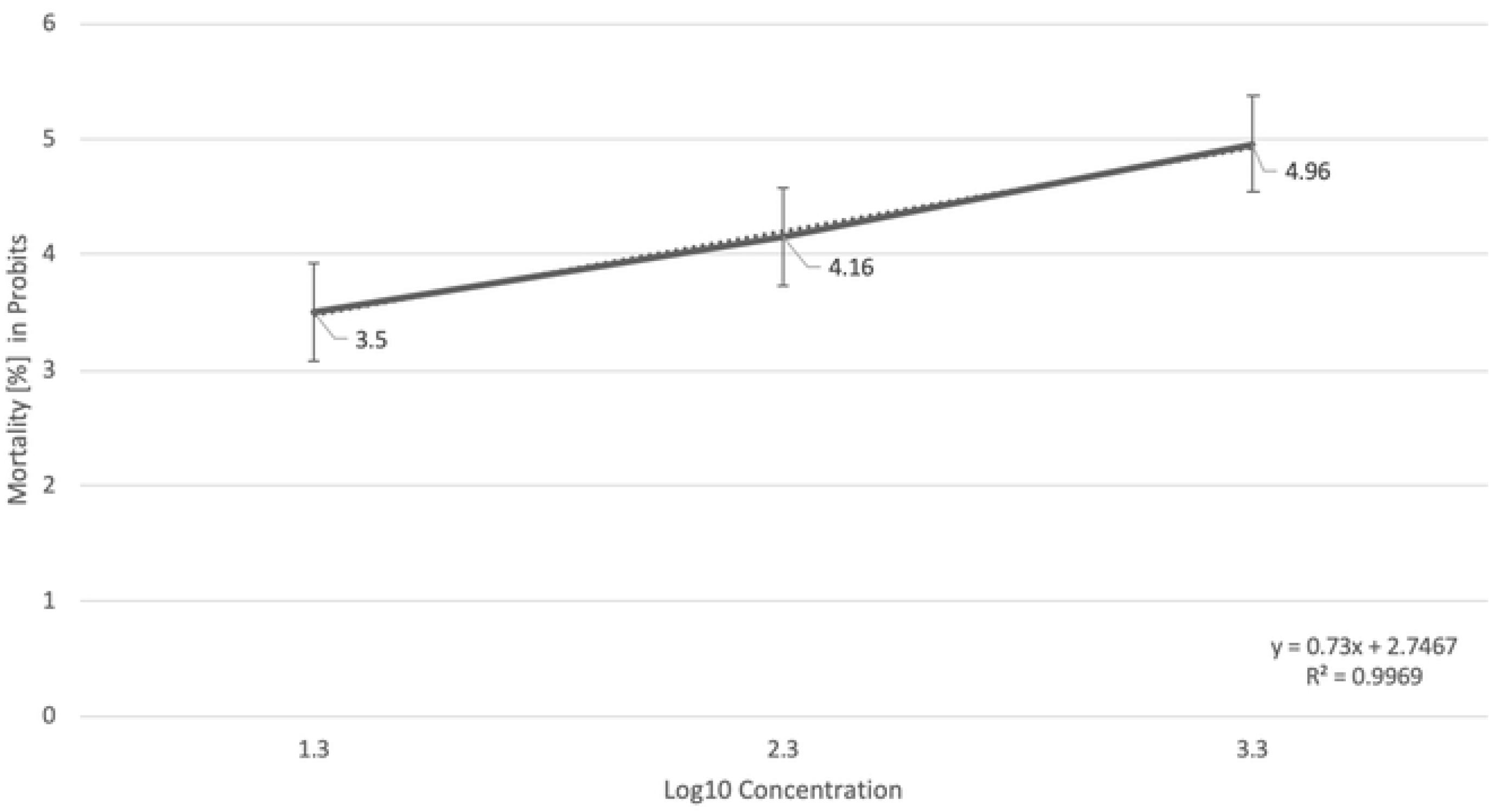
Probit analysis for the estimation of LC_50_ values of *Danio rerio* exposed to PE-MBS. Analyzed results showed that the LC_50_ is 2455.096 μg/L with 95% confidence limits. Error bars indicate standard error.

### Hatching

Hatching is a critical stage in the embryogenesis of *Danio rerio* for it aids in the evaluation of developmental delays and toxicity caused by different substances [93]. A normally developing *Danio rerio* typically hatches between 48 to 72 hpf [94]. *Danio rerio* embryos hatched in the 96 hpf mark are considered late hatchers whereas those hatched in 48 hpf are considered early hatchers [95].

Based on the results shown in Fig 6, the majority of *Danio rerio* embryos hatched at 72 hpf. Embryos treated with 5% ethanol showed the highest number of hatching at 96 hpf. The trend in Fig 6 shows an increasing number of embryos hatching at 48 hpf upon increase in PE-MBS concentration. With these results, administration of statistical analysis of ANOVA indicated that there is a significant difference between the means and variances of the number of hatched *Danio rerio* within the 96 hour exposure to all treatments (S13 Appendix).

**Fig 6.**
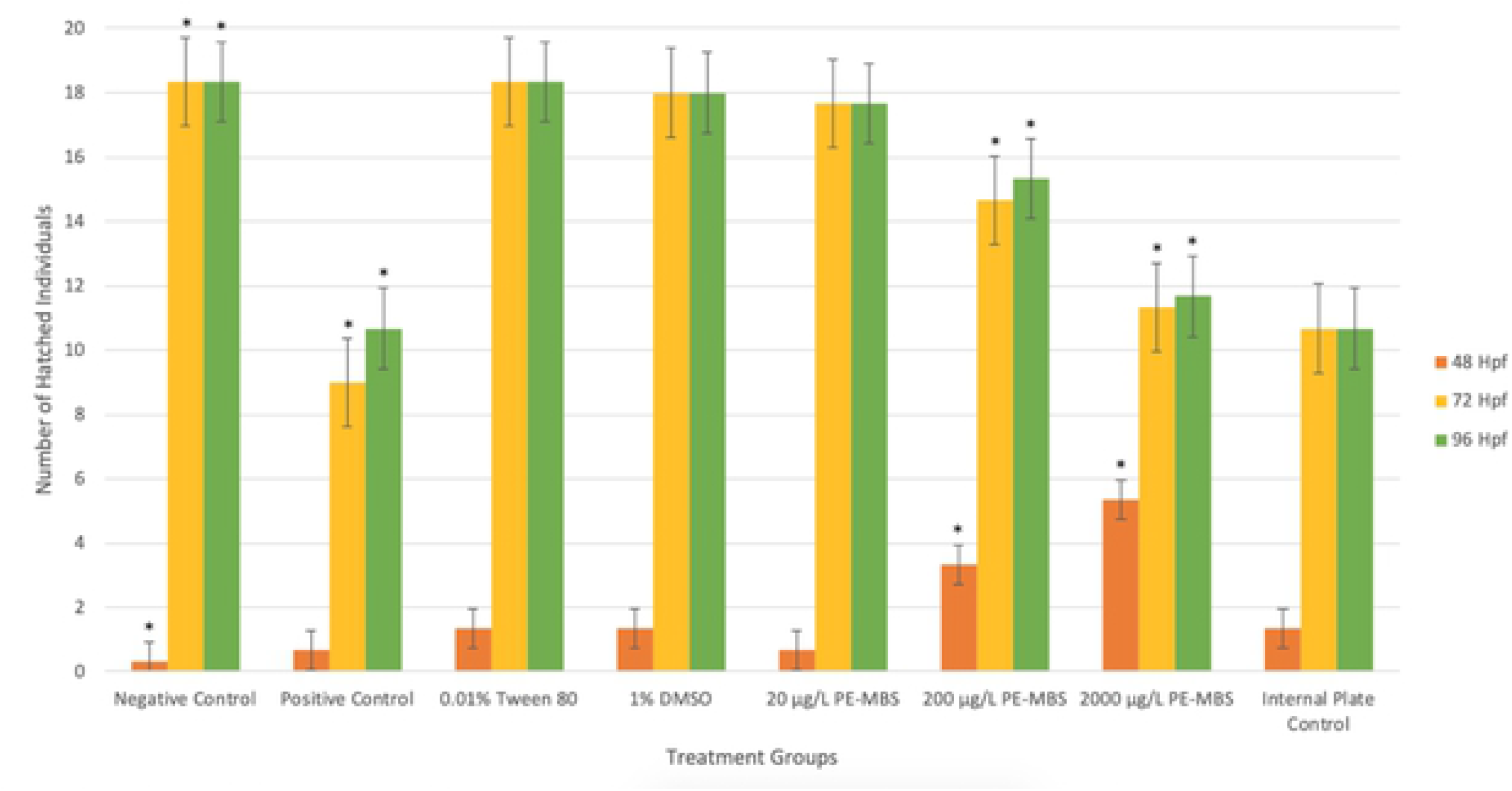
Cumulative number of hatched *Danio rerio* within 96 hour exposure to different treatments. Data shown is based on the average of three replicates performed in the study. Error bars indicate standard error. Single-asterisk indicates a statistically significant difference of cumulative hatching between *Danio rerio* (p< 0.05). (*:p < 0.05).

All throughout the 96 hour exposure, results from Dunnet’s test indicated that no significant difference in the means of the cumulative number of hatched individuals in *Danio rerio* treated with 0.01% Tween 80 and 1% DMSO with the negative control was present (S14 Appendix). Although these substances cause teratogenic and embryotoxic effects in high doses [96,97], they were diluted in accordance to OECD guidelines to induce effects of negligible difference with the negative control while serving as appropriate solvents for the toxicant under study [17].

At 48 hpf, Dunnet’s test revealed that no significant differences were noted in the means of the number of hatched *Danio rerio* treated with 20μg/L PE-MBS and 5% ethanol with the negative control but there is a notable difference for the results garnered for *Danio rerio* treated with 200 and 2000 μg/L PE-MBS with the negative control. This indicates that high concentrations of PE-MBS induces early hatching in *Danio rerio* embryos. This physiological phenomenon may be a result of hypoxia caused by PE-MB. All vertebrates rely on diffusion for both gas exchange and respiratory gas transport, especially in the early stages of development [89]. Microbeads used in the study measured 300-355 μm in size whereas the diameter of the chorionic pores in *Danio rerio* measures less than 1 μm [88]. Possible adherence of PE-MB in the chorionic membrane of *Danio rerio* during embryogenesis may have resulted in clogged pores, hindered gas exchange and consequently, insufficient oxygen supply. As stated in a study by Burrggren and Pinder [89], hypoxia in *Danio rerio* embryos increases truncal muscle movement to agitate water contained inside the chorion and accelerates certain metabolic and respiratory processes to compensate for lack of oxygen [98]. These stress-induced responses as a result of hypoxic environment is also accompanied by premature hatchings since removing the resistance of the chorionic membrane is known to increase oxygen uptake in *Danio rerio* embryos [89]. Notably, recorded data revealed that despite hatching earlier than other treatment groups, *Danio rerio* treated with 200 and 2000 μg/L PE-MBS had lower survival rates after hatching (S4 Appendix). This reinforces the hypothesis that exposure to high doses of PE-MB is both teratogenic and embryotoxic.

For both 72 and 96 hpf, the means of the cumulative number of hatched individuals treated with 20 μg/L PE-MBS did not have a significant difference with the negative control, supporting early speculations that 20 μg/L PE-MBS is not sufficient enough to induce developmental delays nor premature hatching. However, Dunnet’s test results for 5% ethanol garnered a significant difference since a number of embryos died before hatching. The same is true for *Danio rerio* treated with 200 and 2000 μg/L PE-MBS since polyethylene microplastics have the ability to induce embryotoxic effects. Remarkably, a number of late hatchers were noted in *Danio rerio* treated with 5% ethanol as ethanol is known to cause harmful complications and slow down certain processes such as hatching and heart rate [97].

Results from Tukey-Kramer’s test for the cumulative number of hatched individuals within the 96 hour exposure revealed that all three concentrations of PE-MBS are significantly different from each other thus indicating that the rate of premature hatching in *Danio rerio* is dose-dependent and steadily increases depending on the dose administered (S15 Appendix).

### Teratogenicity

Teratogenic endpoints are important to determine the teratogenic potential of a chemical [99] and to generalize the response of *Danio rerio* towards this toxicant [81] of varying concentrations. The most common malformations were edema, bent tail, bent body axis, and collapsed swim bladder. Edema is defined by the accumulation of pellucid fluid in the pericardium or in the yolk sac. A bent tail is observed in an abnormal, dorsoventral or lateral flexion of the tail at the axial level of the caudal fin. A bent body axis is observed in an abnormal flexion of the primary axis. Lastly, a collapsed swim bladder may be more unexpanded than the normal phenotype of a *Danio rerio* swim bladder [96].

As shown in Fig 7, the average number of malformations observed in all treatments and controls were in the following decreasing order: 2000 μg/L PE-MBS, positive control (5% ethanol), 200 μg/L PE-MBS, 20 μg/L PE-MBS whereas 1% DMSO, and 0.01% Tween 80 garnered the same value (S6 Appendix). The embryos in the negative control and the internal plate control did not show any malformations. With these results, administration of statistical analysis of the Kruskal-Wallis Test indicated that there is a significant difference between the means and variances of the number of malformations observed in *Danio rerio* within the 96 hour exposure to all treatments (S16 Appendix). The trend showed that increasing PE-MBS concentration resulted in an increased number of malformations as well.

**Fig 7.**
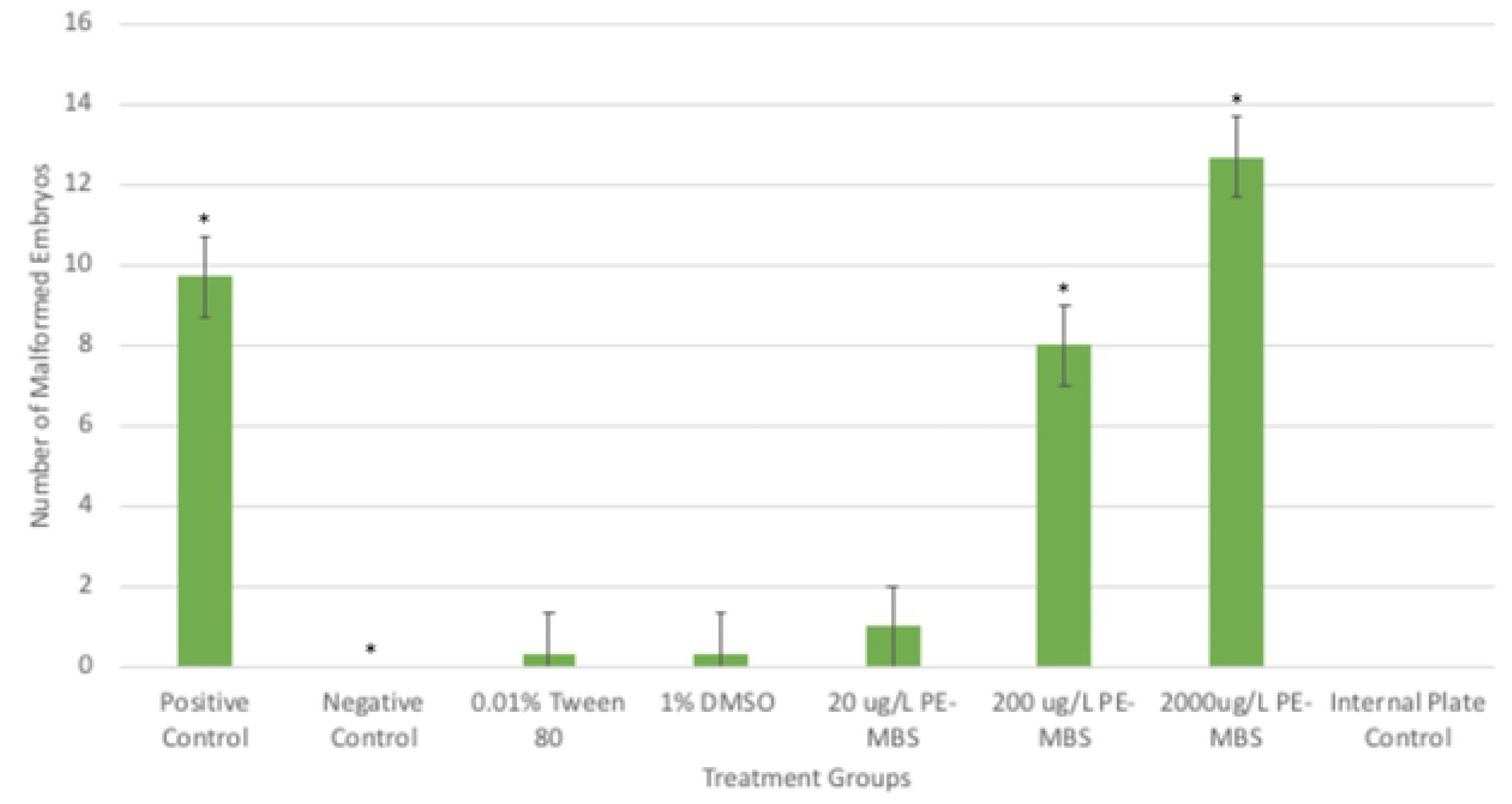
Total number of malformations observed in *Danio rerio* for each treatment at 144 hpf. Error bars indicate standard error. Single-asterisk indicates a statistically significant difference of total number of malformations between *Danio rerio* at 144 hpf (p < 0.05). (*:p< 0.05).

As shown in Fig 8, edema had the highest number of incidents for each group, garnering a range of percent values from 62% to 100%. Bent body axis at 8% to 33% and bent tail with percent values of 21% to 24% came next while the collapsed swim bladder was the least observed teratogenic endpoint for all groups, garnering percent values from 3% to 13%.

**Fig 8.**
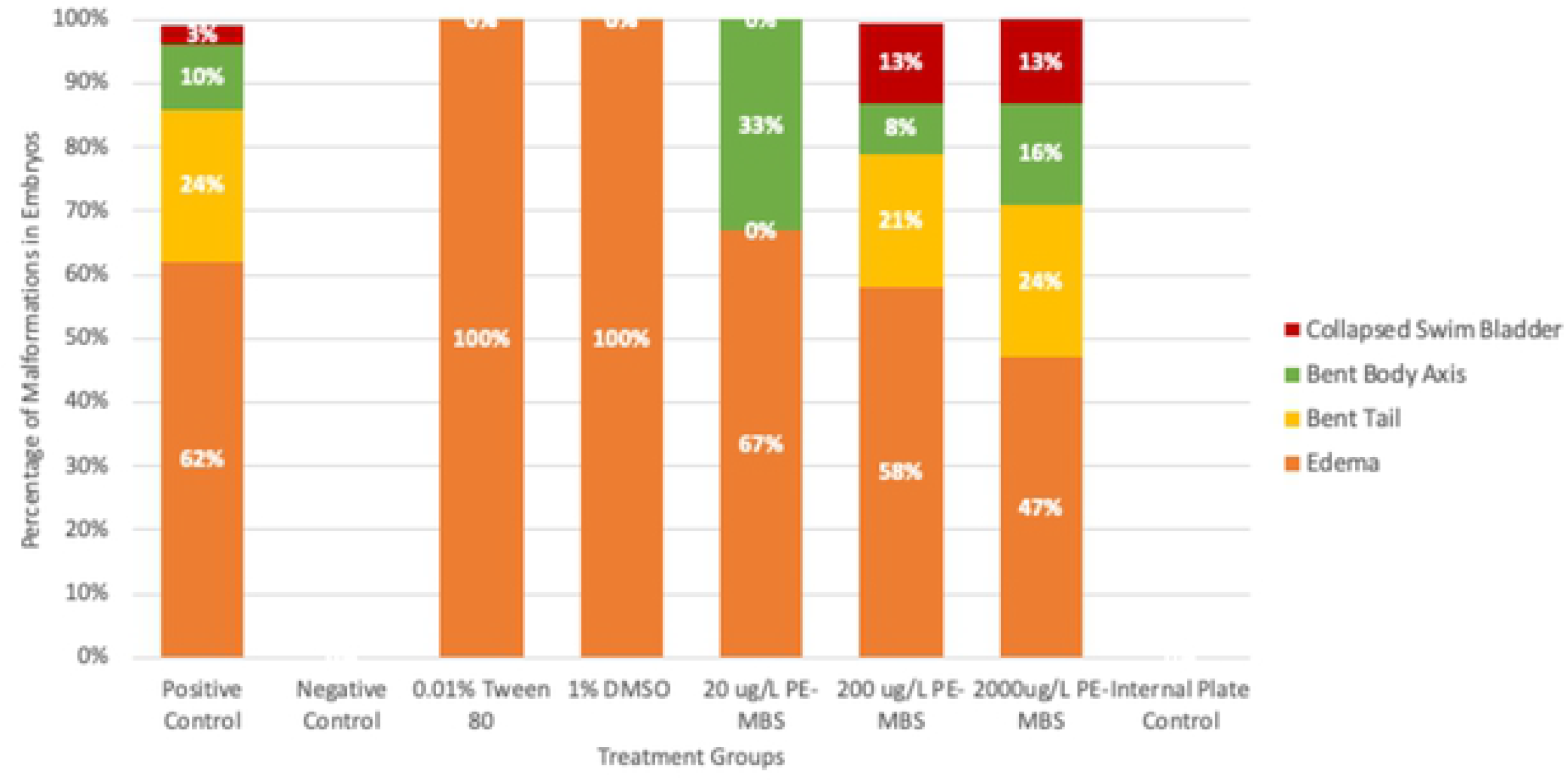
Relative percentages of malformations observed in *Danio rerio* for each treatment at 144 hpf. Percentage shown is based on the average of three replicates performed in the study.

According to Ali et al. [96], one of the abnormalities found in *Danio rerio* subjected to 8000 mg/L ethanol was pericardial edema. Another study [100] found that even at lesser concentrations of ethanol (1.5% and 2%), abnormalities such as bent body axis were observed in *Danio rerio* embryos. That being said, all literature coincided with the observations in embryos treated with 5% ethanol in this study (Fig. 9B, 10C, 11A, 11B). Dunn’s Test (S17 Appendix) had also indicated that the positive control, the 5% ethanol significantly differed with the negative control.

**Fig 9.**
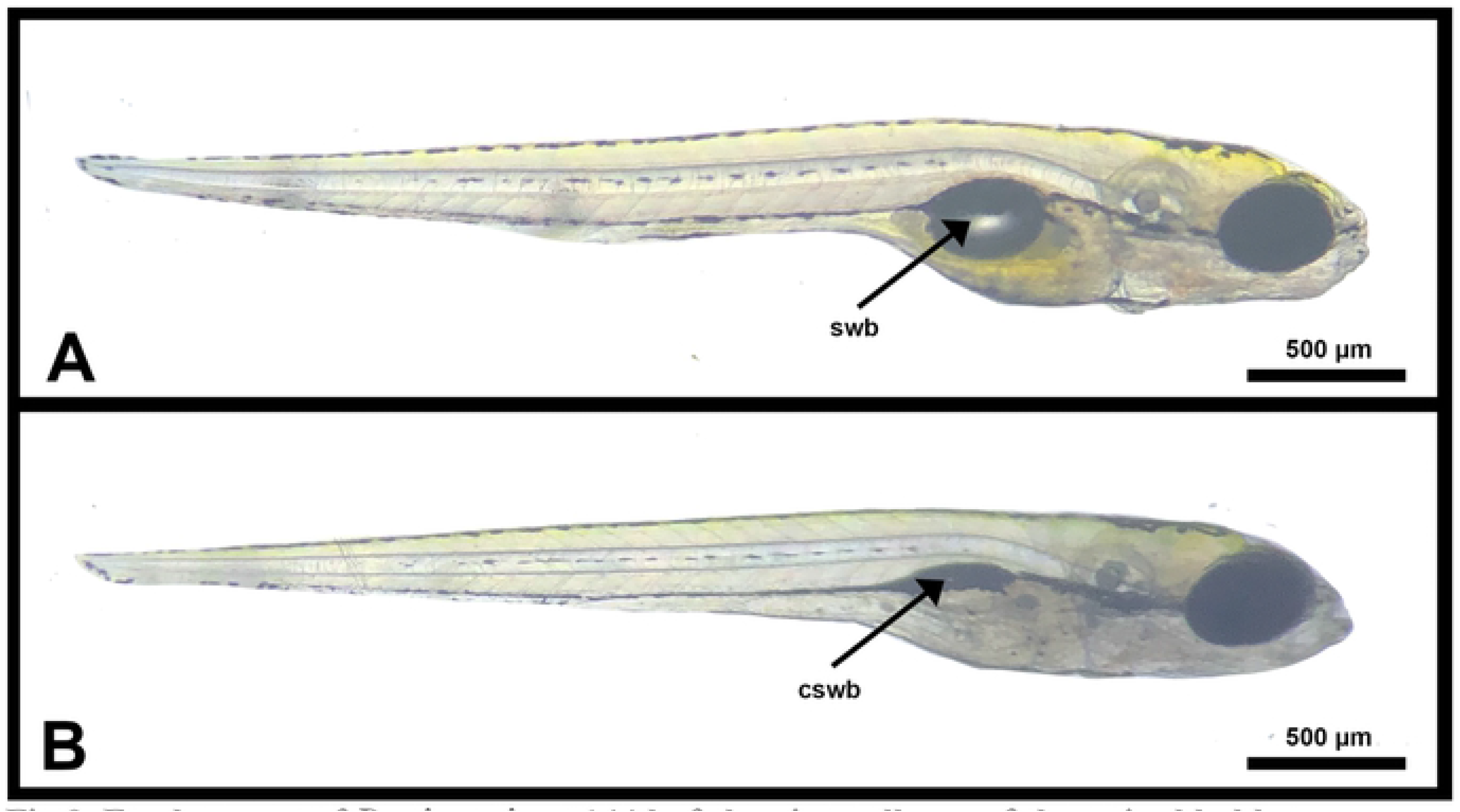
Fresh mount of *Danio rerio* at 144 hpf showing collapse of the swim bladder. (A) shows normal development of the swim bladder (swb) observed in the negative control (RW), 0.01% Tween 80 % DMSO, and 20 μg/L PE-MBS. (B) shows collapsed swim bladder (cswb) observed in the following treatments of increasing order: the positive control (5% ethanol), 200 μg/L PE-MBS, and 2000 μg/L PE-MBS.

**Fig 10.**
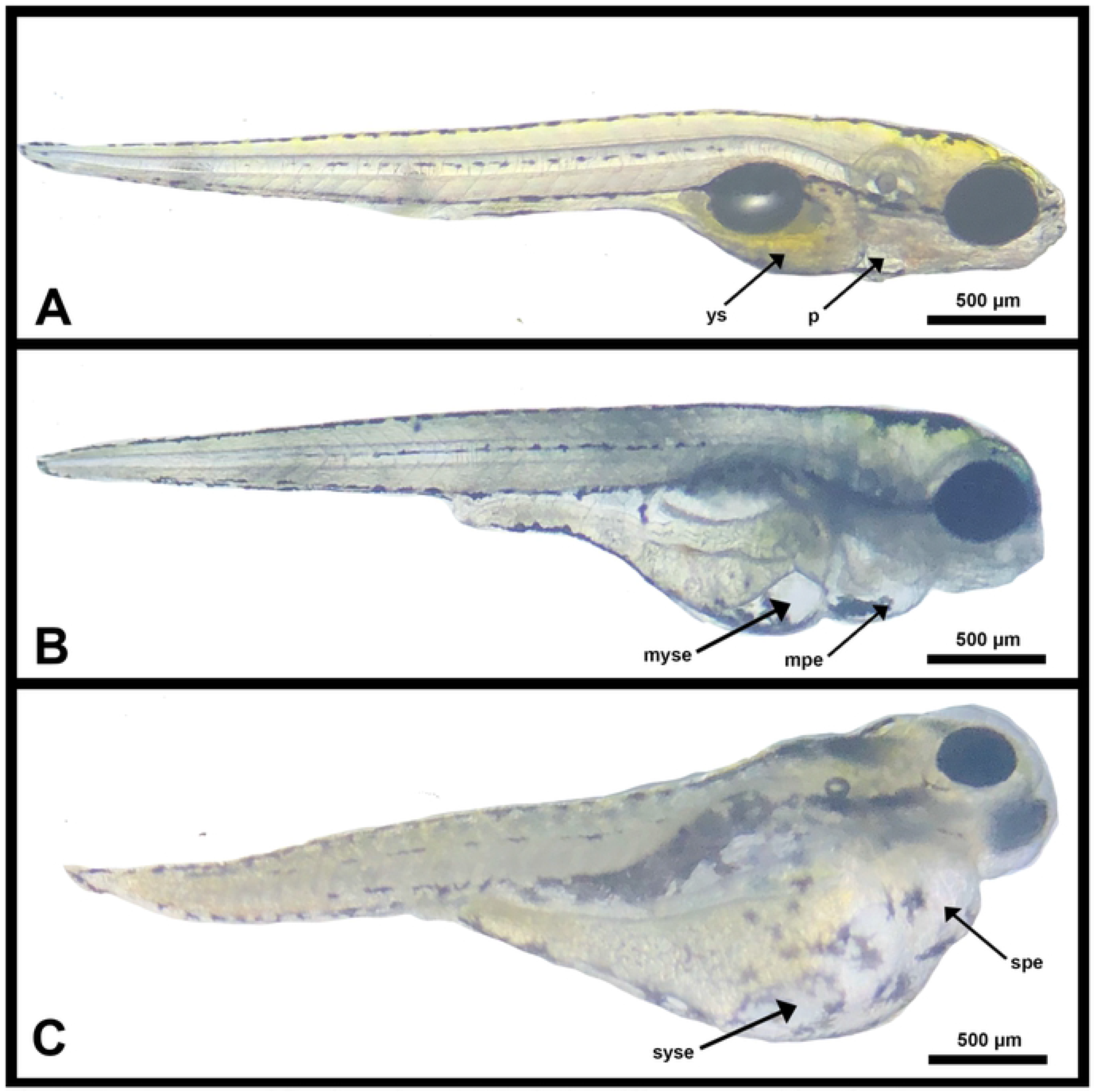
Fresh mount of *Danio rerio* at 144 hpf with different severities of yolk sac and pericardial edema. (A) shows the normal development observed in the negative control (RW) with normal yolk sac (ys) and pericardium (p). (B) exhibits mild yolk sac edema (myse) observed in treatments 0.01% Tween 80 and 1% DMSO and mild pericardial edema (mpe). (C) shows severe yolk sac (syce) and pericardial edema (spe) both observed in 200μg/L PE-MBS, 2000μg/L PE-MBS, and 5% ethanol with the last two respective treatment and control groups exhibiting the most incidents of edema.

**Fig 11.**
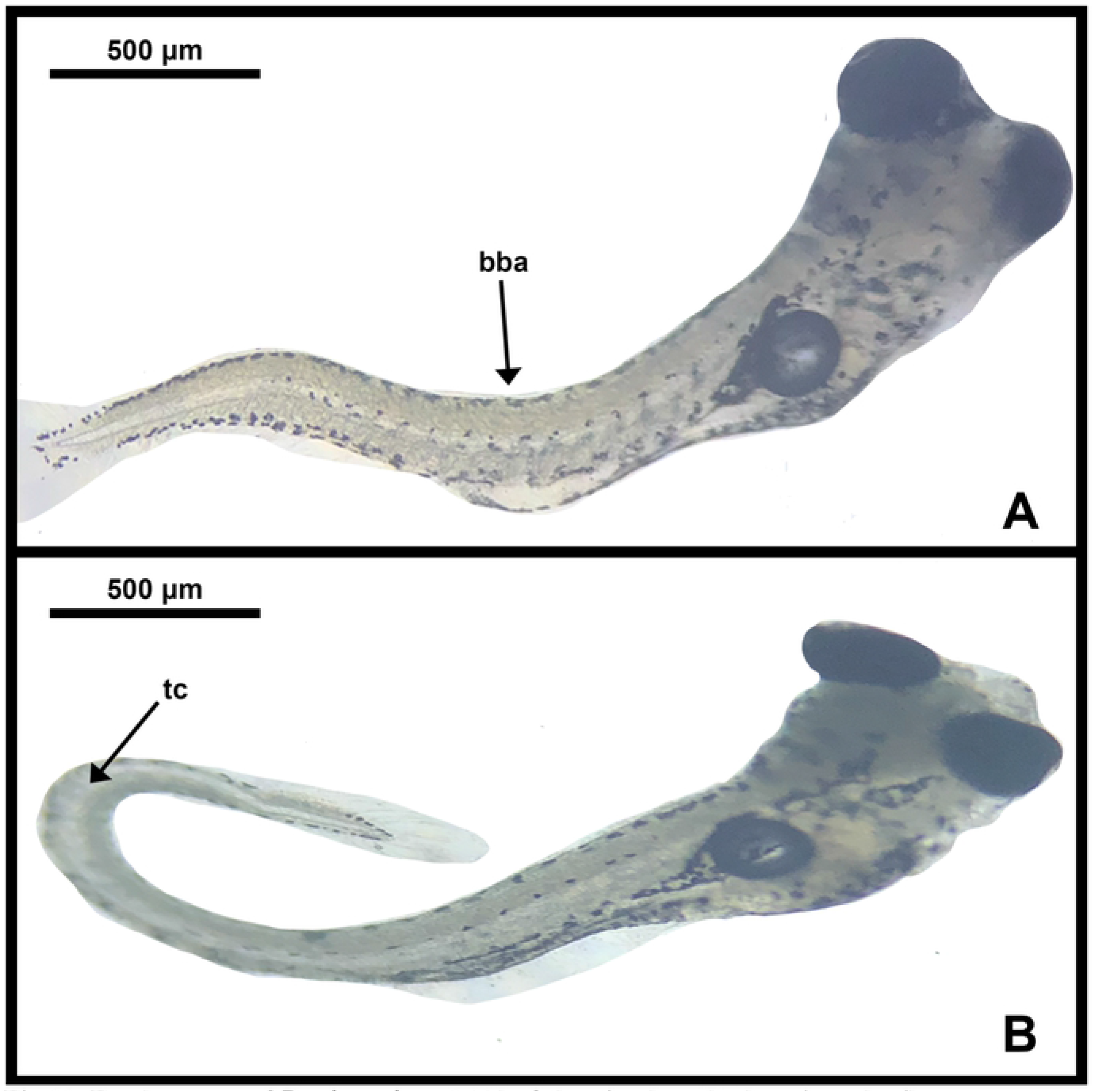
Fresh mount of *Danio rerio* at 144 hpf showing bent body axis and tail curvature. (A) exhibits bent body axis (bba). (B) exhibits tail curvature (tc). Both malformations were observed in the positive control (5% ethanol) and in treatments, 200 and 2000 μg/L PE-MBS. Bent body axis was observed in the 20 μg/L PE-MBS.

Ali et al. [96] stated that *Danio rerio* embryos subjected to 200 mg/L Tween 80 exhibited dispersed pigment cells, bent body axis, and branchial arch hypoplasia. However, since Tween 80 was diluted to lesser concentrations in this study, the mean observation found in *Danio rerio* embryos subjected to the resulting concentration was too negligible to significantly differ with the negative control (Fig. 9A, 10B, 11A). Meanwhile, DMSO was reported as a teratogen at higher concentrations [101]. However, at lesser concentrations likened to 1%, embryos treated with 1% DMSO did not exhibit significant teratogenicity (Fig. 9A, 10B, 11A). This finding also coincided with other studies [101,81]. Statistically, 0.01% Tween 80 and 1% DMSO both did not significantly differ with the negative control as well (S17 Appendix).

Two hundred (200) and 2000 μg/L PE-MBS treatment groups significantly differed with the negative control (S17 Appendix); however, concentration of 20 μg/L PE-MBS treatment was too negligible to significantly differ with the negative control. With that said, polyethylene microbeads may affect the body axis, body proportion and other morphological parameters of aquatic organisms depending on PE-MBS concentration. Their sizes [102] may also be a factor associated with malformation.

The Tukey Kramer Test indicated significant differences between all PE-MBS treatment groups. Results also showed that the number of deformities increased upon increase of PE-MBS concentration. This may be interpreted that higher concentration of PE-MBS induces greater teratogenicity in *Danio rerio*. In spite of early speculation, polyethylene microbeads may have caused disturbance to regulating barriers in internal water diffusion [5], possibly substantiating increasing incidents of edema in increasing concentrations. Edema being the highest number of type of malformation in all PE-MBS treatment groups, may be regarded as a symptom of hypoxia in *Danio rerio* embryos, further substantiating that PE-MB may cause hypoxia [5]. It has also been observed that sublethal stages of hypoxia can increase embryonic fish malformations by 77.4% ultimately resulting in decline of species’ fitness and aquatic populations [103].

Bent tails have also been reported in microplastic-treated *Danio rerio* adults at moderate and high concentrations [104], coinciding with another article as well [105]. In the former study, bent tails observed in polyethylene microplastic treated *Danio rerio* may be associated with “knockdown of the cysteine-rich motor neuron 1 gene (crim1) or missense mutation in polycystin-2(pkd2).” This gene encodes for the activation of the Ca^2+^ cation channel which is important in the skeletal muscle excitation-contraction [106] that may be depicted in the tail movement of the zebrafish. However, Kaleuff [107] recommends further investigation regarding whether exposure to microplastics significantly changes the level of target gene expression and phenotype. Bent body axes were also observed in polyethylene microplastic treated *Danio rerio* embryos [5]. Further observation and findings are needed to associate these teratogenic effects to adhesion of polyethylene microbeads to the external surface and to the gastrointestinal system of the *Danio rerio* embryo; however a study of Malafaia et al. [5] associates these teratogenic effects to this occurrence.

Collapsed swim bladders have also been evident in the 200 μg/L PE-MBS treatment but more especially in the 2000 μg/L PE-MBS treatment (Fig. 9B). The swim bladder, an aid in making upward hydrodynamic forces in prevention from sinking [108], was exhibited to be collapsed in *Danio rerio*, and this may have resulted from hypoxia [107] possibly induced by 200 μg/L and 2000 μg/L PE-MBS. Collapsed swim bladders have also been observed in *Danio rerio* embryos affected by nano plastics [11].

With substantiating the results from the Tukey Kramer Test indicating significant differences between all PE-MBS treatment groups (S19 Appendix), it can be said that teratogenicity increases with increasing PE-MBS concentration.

### Heartbeat

The heart rate of a developing zebrafish embryo is usually at 120-180 beats per minute (bpm) and it is usually visible at 48 hpf. It is a significant endpoint that should be observed to ensure tissue perfusion in all parts of the developing embryo [109].

As shown in Fig 12, the general trend was that the fastest bpm was always evident in the 96 hpf while the slowest bpm was observed in the 48 hpf for each treatment and control group. This is because the heart rate increases as development takes place [109]. As shown in the Fig 12, embryos in PE-MBS treatments steadily increased heart rate as the PE-MBS concentration increased. As for the control groups, each group showed steady increase of heart rate in exception to the positive control. Embryos in the positive control showed a slower rate of heartbeat as each succeeding 24th hour went by.

**S12 Fig.**
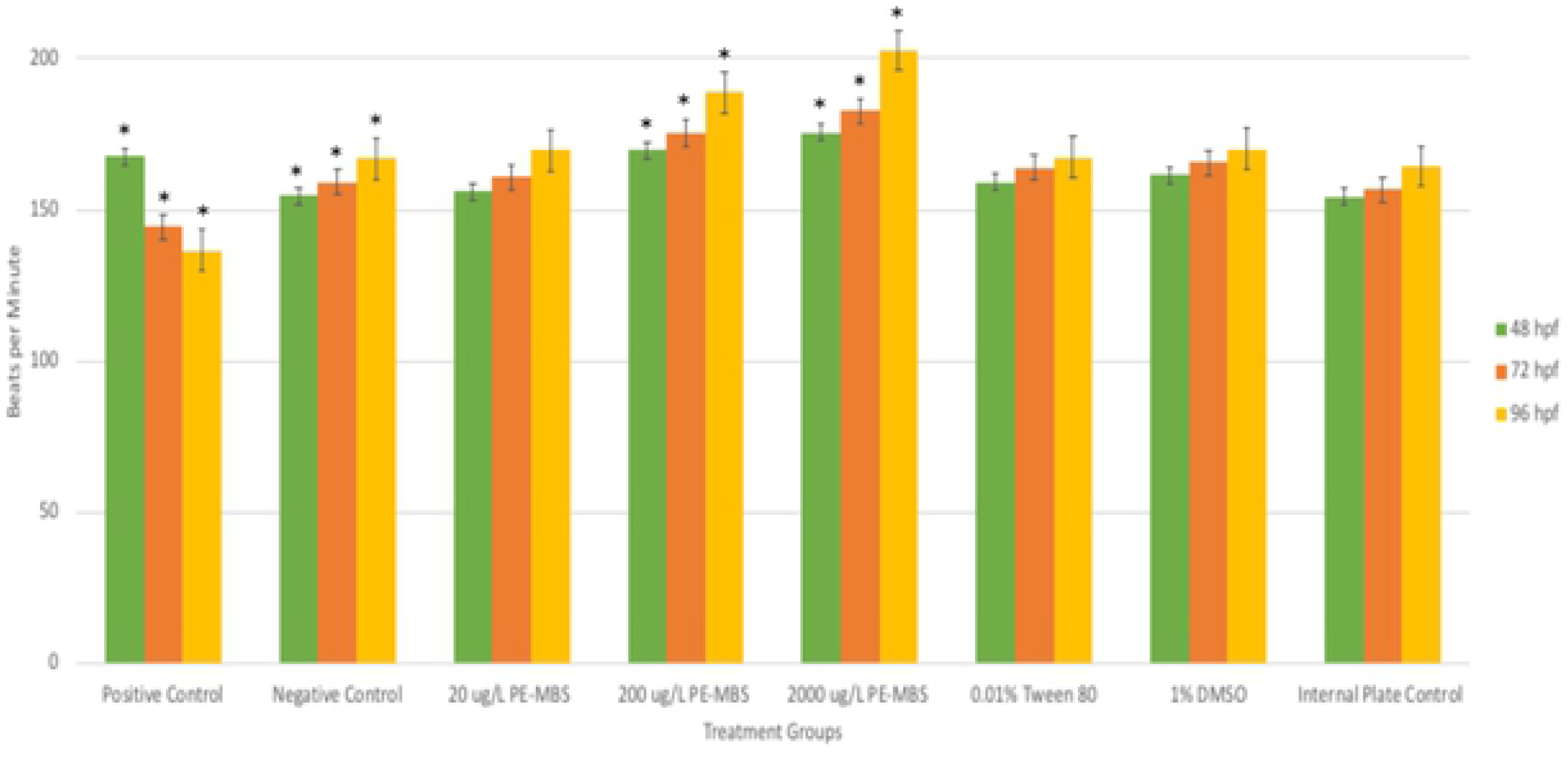
Heart rate (bpm) observed in *Danio rerio* for each treatment. Data shown is based on the average of three replicates performed in the study. Error bars indicate standard error. Single-asterisk indicates a statistically significant difference of heart rate between *Danio rerio* (p< 0.05). (*:p< 0.05).

But among all groups, the fastest average bpm was evident in embryos treated with 2000 μg/L PE-MBS at 96 hpf (i.e., 203 bpm) while the slowest average bpm was observed in the positive control at 96 hpf (i.e., 137 bpm).

With these results, statistical analyses of ANOVA indicated significant differences between the means and the variances of each treatment (S20 Appendix). These results led to *post hoc* analysis, namely the Dunnet’s Test (S21 Appendix) and the Tukey Kramer Test (S22 Appendix). At all hpf, means of all treatments had significant differences with those in the negative control in exception to ones in 0.01% Tween 80, 1% DMSO and 20 μg/L PE-MBS. Meanwhile, the latter test indicated significant differences between all PE-MBS treatments.

In a normal embryonic development of *Danio rerio*, the heart rate increases as development takes place [109] as earlier mentioned. Increased heart rate had been observed in all treatments in exception to 5% ethanol. Ethanol may cause decrease in size of ventricles and lessen the number of cardiomyocytes in the heart of a developing zebrafish [110], leading to mortality and effects of teratogenicity. Hallare et al. [81] also stated that ethanol greater than 1.5% concentration caused developmental delays in heart beating. This finding also coincided with other studies [80,111]. Hence, 5% ethanol had a significant difference with the negative control for all hpf.

For other chemicals, means of heart rate in 0.01% Tween 80 and in 1% DMSO did not significantly differ with the negative control. Tween 80 is known to have a relatively high toxicity towards *Danio rerio,* even more than its toxicity to rodents because of its surfactant properties [76]. But in comparison to this study, Tween 80 was diluted to a lesser concentration in accordance with OECD guidelines [17] hence its concentration was found to be too negligible to cause a significant difference with the negative control. High concentrations of DMSO (≥ 1.5% v/v) were observed to induce brachycardia and pronounced arrhythmia; but at lower concentrations, initial increase of average heart rate was observed instead [81].

20 μg/L PE-MBS did not significantly differ with those in the negative control. This may be interpreted that 20 μg/L PE-MBS was an insufficient dose that could not induce an irregularly increased bpm in comparison to the bpm observed in the negative control; however, the other higher PE-MBS concentrations induced a significantly increased heart rate in comparison to the negative control that also exhibited an increase of bpm but at a steady rate. Results may be interpreted that the higher the PE-MBS concentration, the more likely *Danio rerio* will be subjected to cardiac toxicity. It is said that exposure to polyethylene causes cardiac toxicity, a term defined by a greatly increased heart rate that may be attributed to physiological stress [112].

This significant increase of heart rate may also be associated with acute hypoxia contributed by adherence of polyethylene microbeads to chorionic membranes as earlier stated. One of the symptoms of hypoxia may be irregular, rapid heartbeat [113]. The study of Crail [112] also showed significant increase of heart rate at delimited oxygen concentration. That being said, means observed in 200 μg/L PE-MBS at 72 and 96 hpf and means observed in 200 μg/L PE-MBS at 96 hpf have exceeded the regular heart rate of a *Danio rerio* embryo. This further substantiates the occurrence of irregular, rapid heart rate in the zebrafish embryo at increasing concentrations and at longer exposure to PE-MBS.

Tukey Kramer *post hoc* test (S22 Appendix) revealed that the heart rate of *Danio rerio* treated with different concentrations of PE-MBS significantly differed from each other at all hpf. Heart rate of zebrafish embryos increased and became more irregular upon exposure to increasing concentrations of PE-MBS, implying that cardiac toxicity due to PE-MBS may be dose dependent.

### Conclusions and recommendations

The inclusion of polyethylene microbeads in personal care products such as facial washes and cosmetics has captured the attention of the scientific community due to the deleterious effects it poses on aquatic organisms. In this study, *Danio rerio* was chosen to be the representative model due to its availability, high fecundity and great similarity with the human genome. Polyethylene microbeads used in the study were based on measurements similar to actual commercial facial cleansers that contained polyethylene. Results from the Fish Embryo Acute Toxicity Test revealed that 20 μg/L did not have significant difference with the negative control in the observed parameters (i.e., embryotoxicity, teratogenicity, hatching, and heartbeat), but 200 and 2000 μg/L did, ascertaining that static exposure to high concentrations of polyethylene microbeads is embryotoxic and teratogenic to *Danio rerio* embryos. Cases of mortality may be due to the soft extraction of polyethylene microbeads using 1% DMSO that may have induced leaching of toxic additives and Endocrine-Disrupting Chemicals (EDCs). In accordance with literature outside of this study, these leached chemicals may have disrupted metabolic pathways [5,85], alter gene integrity [85,86], and cause cell apoptosis [87]; hence possibly resulting in *Danio rerio* toxicity during embryogenesis; however, this needs a more thorough study at gene level. Adherence of polyethylene microbeads to the chorionic membrane may also have disrupted gas exchange and induced hypoxia [5]. Hypoxia may have been the underlying cause of observed premature hatching in *Danio rerio* which, in effect, increased occurrences of larval death and incidences of teratogenic abnormalities such as edema, collapsed swim bladder, and bent body axes. Rapid and irregular heart rate was also observed among *Danio rerio* embryos and may be associated with acute hypoxia and cardiac toxicity caused by polyethylene microbead exposure.

The result obtained from the computation for the LC_50_ is 2455.096 μg/L and is higher than the treatment concentrations used in the study. Nonetheless, strong and urgent actions against the production of facial cleansers containing PE-MB must be implemented to reduce the microplastic pollution in bodies of water. Furthermore, investments and policy reforms on improving plastic wastes management must also be enacted to minimize microplastic leaching into the aquatic ecosystem from wastewater treatment plants. Through these concerted efforts, issues of bioaccumulation and toxicity by microplastic (e.g., microbeads) pollutants may be mitigated or prevented and consequently save the lives of both aquatic organisms and humans alike.

Since this study was only limited to polyethylene, different types of microbeads such as polypropylene and polyamide and different sizes ranging less than 300-355 μm may be included to broaden the study. It is also recommended that zebrafish exposed to microbeads may be further observed at the gene level to investigate the possible occurrence of mutations and other alterations such as hsp70, crim1, and pkd2 [105] that contribute to embryotoxicity and teratogenicity of the zebrafish. New biomarkers can also be searched further that can be used to monitor the health of aquatic habitat and its biota.

## Acknowledgments

This study was made possible from the grant given by University of the Philippines National Institute of Health and the laboratory facilities UP Manila College of Arts and Sciences have provided to conduct the proper feeding and maintenance of the zebrafish. In lieu of its feeding and maintenance, it is with deepest gratitude that the researchers reach out to Sir Edgar Acantilado and Sir Maxcitar Amar for helping them with the process of the experimentation from the beginning until the end. The researchers would also like to thank Ma’am Julieta Dator Holasca from Central Luzon State University for her kind accommodation when the researchers purchased the zebrafish used in this study. Finally, the researchers would also like to extend their deepest gratitude to Ma’am Margaret L.C. De Guzman for her constant guidance and encouragement all throughout the duration of this study. Significant contributions and comments put in earlier versions of the thesis manuscript given by Ma’am Margaret L.C. De Guzman, Sir Arnold V. Hallare, and Sir Jay T. Dalet have been more than helpful in making the study more productive and more apt enough for the next researchers who want to continue this study.

## Supporting Information

**S1 Table. Experimental setup for the Fish Embryo Acute Toxicity Test.**

**S1 Appendix. Number of fertilized and unfertilized eggs collected during spawning.**

**S2 Appendix. Toxicological endpoints observed in *Danio rerio* for each trial within the 96 hour exposure.**

**S3 Appendix. Cumulative number of deceased *Danio rerio* for each trial within the 96 hour exposure.**

**S4 Appendix. Cumulative number of deceased *Danio rerio* embryo and larvae for each trial at the end of the-96 hour exposure.**

**S5 Appendix. Cumulative number of hatched *Danio rerio* for each trial within the 96 hour exposure.**

**S6 Appendix. Average number of malformations observed in Danio rerio embryos for each trial at 144 hpf.**

**S7 Appendix. Average number of each kind of malformation observed in *Danio rerio* for each trial at 144 hpf.**

**S8 Appendix. Average bpm of *Danio rerio* embryos exposed for each trial within the 96 hour exposure.**

**S9 Appendix. Single factor analysis of variance for the effect of different treatments to the mortality of *Danio rerio* within the 96 hour exposure.**

**S10 Appendix. Dunnet’s test for cumulative mortality of *Danio rerio* exposed to different treatments within the 96 hour exposure with 95% confidence intervals.**

**S11 Appendix. Tukey HSD/Kramer test for cumulative mortality of *Danio rerio* treated with varying concentrations of PE-MBS within the 96 hour exposure.**

**S12 Appendix. Calculation of LC_50_ of PE-MB using probit analysis with 95% confidence limits**

**S13 Appendix. Single factor analysis of variance for the effect of different treatments to the cumulative hatching of *Danio rerio* within the 96 hour exposure.**

**S14 Appendix. Dunnet’s test for cumulative hatching of *Danio rerio* exposed to different treatments within the 96 hour exposure with 95% confidence intervals.**

**S15 Appendix. Tukey HSD/Kramer test for cumulative hatching of *Danio rerio* treated with varying concentrations of PE-MBS within the 96 hour exposure.**

**S16 Appendix. Kruskal-Wallis Test for the malformations observed in *Danio rerio* exposed to different treatments at 144 hpf.**

**S17 Appendix. Dunn’s test for malformations observed in *Danio rerio* embryos at 144 hpf to different treatments (p<0.05).**

**S18 Appendix. Single factor analysis of variance for the effect of varying concentrations of PE-MBS to the number of malformations observed in *Danio rerio* at 144 hpf.**

**S19 Appendix. Tukey HSD/Kramer test for malformations of *Danio rerio* treated with varying concentrations of PE-MBS at 144 hpf.**

**S20 Appendix. Single factor analysis of variance for the effect of different treatments to the heart rate (bpm) of *Danio rerio* within the 96 hour exposure.**

**S21 Appendix. Dunnet’s test for the heart rate (bpm) in *Danio rerio* exposed to different treatments within the 96 hour exposure with 95% confidence intervals.**

**S22 Appendix. Tukey HSD/Kramer test for the heart rate (bpm) of *Danio rerio* treated with varying concentrations of PE-MBS within the 96 hour exposure.**

**S23 Appendix. Institutional Animal Care and Use Committee Letter of Approval.**

